# Chemogenetic ligands for translational neurotheranostics

**DOI:** 10.1101/487637

**Authors:** Jordi Bonaventura, Mark A. Eldridge, Feng Hu, Juan L. Gomez, Marta Sanchez-Soto, Ara M. Abramyan, Sherry Lam, Matthew Boehm, Christina Ruiz, Mitchell Farrell, Andrea Moreno, Islam Mustafa Galal Faress, Niels Andersen, John Y. Lin, Ruin Moaddel, Patrick Morris, Lei Shi, David R. Sibley, Stephen V. Mahler, Sadegh Nabavi, Martin G. Pomper, Antonello Bonci, Andrew G. Horti, Barry J. Richmond, Michael Michaelides

## Abstract

Designer Receptors Exclusively Activated by Designer Drugs (DREADDs) are a popular chemogenetic technology for manipulation of neuronal activity in uninstrumented awake animals with potential for precision medicine-based clinical theranostics. DREADD ligands developed to date are not appropriate for such translational applications. The prototypical DREADD agonist clozapine N-oxide (CNO) lacks brain entry and converts to clozapine. The second-generation DREADD agonist, Compound 21 (C21), was developed to overcome these limitations. We found that C21 has low brain penetrance, weak affinity, and low *in vivo* DREADD occupancy. To address these drawbacks, we developed two new DREADD agonists, JHU37152 and JHU37160, and the first dedicated positron emission tomography (PET) DREADD radiotracer, [^18^F]JHU37107. JHU37152 and JHU37160 exhibit high *in vivo* DREADD potency. [^18^F]JHU37107 combined with PET allows for DREADD detection in locally-targeted neurons and at their long-range projections, enabling for the first time, noninvasive and longitudinal neuronal projection mapping and potential for neurotheranostic applications.

---

Precision medicine exhibits considerable advantages over conventional medical treatment. Theranostics is a field within precision medicine that provides safer, better-targeted and more efficacious treatment strategies compared to conventional medicine. By combining therapeutic and diagnostic (i.e. theranostic) practices, we could establish a personalized treatment approach encompassing disease diagnosis, drug delivery, and disease monitoring. Currently, theranostic interventions are mainly utilized in fields like oncology, but in the future, this approach could also revolutionize treatments of brain diseases. Such *neurotheranostic* interventions are particularly timely given recent neurotechnological developments and resulting precise knowledge of neural circuit involvement in neurological and psychiatric diseases.

Designer Receptors Exclusively Activated by Designer Drugs (DREADD)^1^ technology is a powerful chemogenetic approach used for neuromodulation in uninstrumented animals. Combining a DREADD with translational molecular imaging methods such as positron emission tomography (PET) extends this promise to encompass neurotheranostics, because non-invasive monitoring is essential to confirm receptor expression and function. DREADD ligands developed to date are not suitable for central nervous system (CNS) applications^2^. The prototypical DREADD agonist clozapine *N*-oxide (CNO) lacks brain penetrance and metabolizes to the antipsychotic clozapine which is the active *in vivo* CNS DREADD agonist^2^. Therefore new, potent DREADD agonists and selective, high-affinity DREADD PET radioligands are needed to advance the translational potential of this powerful chemogenetic technology.

Recently, a new second-generation DREADD ligand, Compound 21 (C21), was put forward as an effective DREADD agonist with excellent brain penetrance that does not convert to clozapine^3,4^. To determine whether C21 is suitable for activating DREADDs, a series of studies in rodents and nonhuman primates (NHPs) characterizing its activity *in vivo* was undertaken. We first assessed C21 and clozapine (**Fig. 1A**) binding in mouse and rhesus monkey (*Macaca mulatta*) brain sections expressing DREADDs. Low concentrations of [^3^H]clozapine bound to DREADDs in intact tissue sections in both species but [^3^H]C21 did not show any DREADD binding in the mouse (**Fig. 1B**) and showed only weak DREADD binding coupled with high off-target binding in the macaque (**Fig. 1C**). In DREADD-expressing HEK-293 membranes C21 inhibited [^3^H]clozapine binding with low affinity (^hM3Dq^*K*_i_=230 nM; ^hM4Di^*K*_i_=91 nM) compared to clozapine (^hM3Dq^*K*_i_=3.5 nM; ^hM4Di^*K*_i_=2.8 nM) (**Fig. 1D**). In functional assays measuring excitatory hM3Dq DREADD activation in transfected HEK-293 cells, C21 elicited potent intracellular Ca^2+^ increases comparable to that of clozapine (**Fig. 1E**). However, C21 was less potent than clozapine at Gi/o activation in cells or tissue expressing hM4Di (**Figs. 1E-G**). Neither clozapine nor C21 produced detectable functional responses in cells (**Extended Fig. 1**) or tissues devoid of DREADD expression (**Figs. 1F-G**).

**Figure 1.**
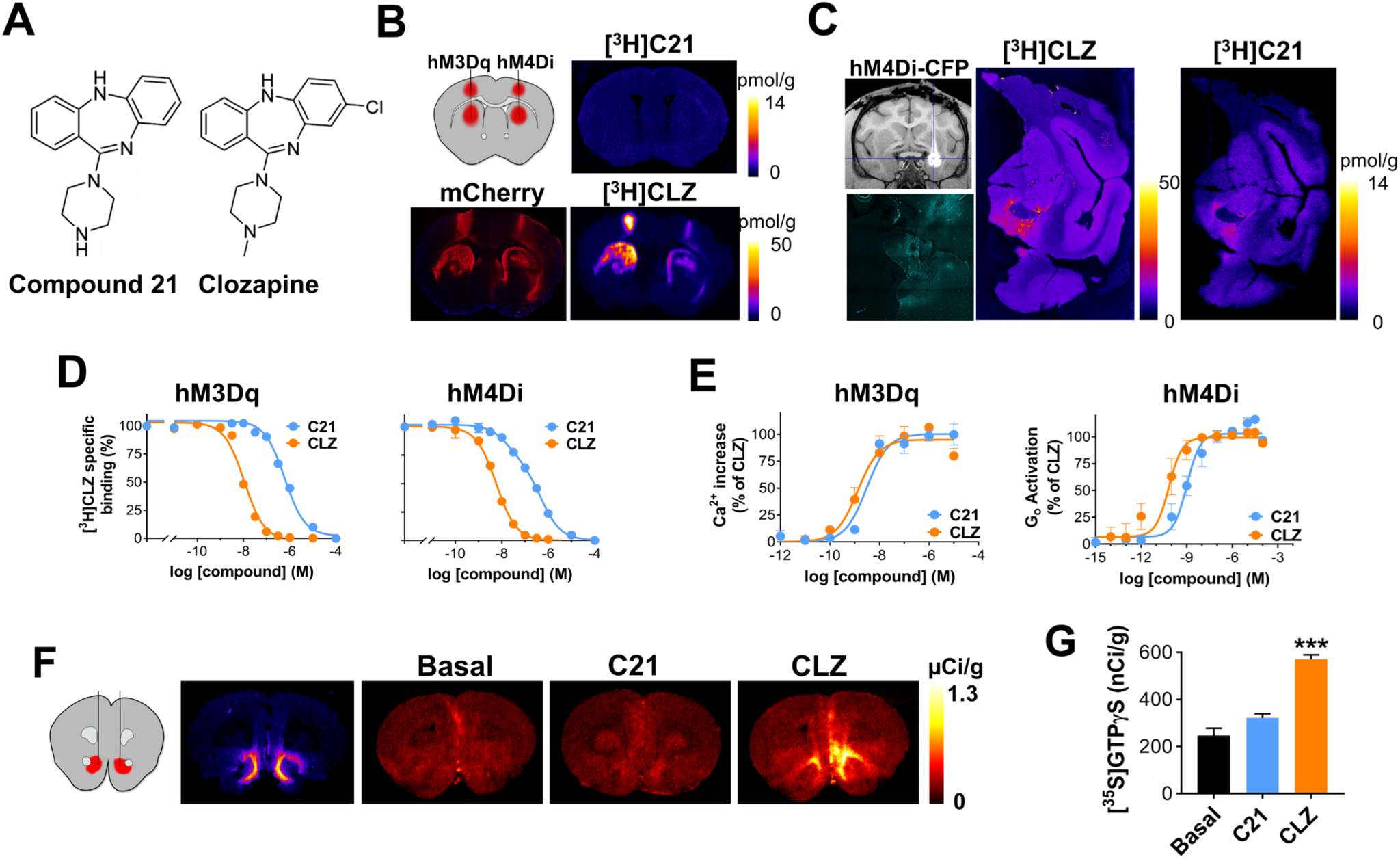
The second-generation DREADD agonist C21 exhibits low *in vitro* DREADD affinity and potency compared to clozapine. (**A**) Structures of C21 and clozapine (CLZ). (**B**) Unlike [^3^H]CLZ, [^3^H]C21 does not bind to DREADDs expressed in mouse brain tissue. (**C**) [^3^H]C21 and [^3^H]CLZ both bind to hM4Di expressed in the monkey amygdala but [^3^H]C21 exhibits lower selectivity. (**D**) C21 displaces [^3^H]CLZ with low affinity whereas CLZ displaces [^3^H]CLZ with high affinity. (**E**) C21 and CLZ exhibit a comparable efficacy profile in cells-expressing hM3Dq but CLZ is more efficacious in cells expressing hM4Di. (**F, G**) CLZ induces significantly greater hM4Di activation ([^35^S]GTP*γ*S binding) compared to C21 in rat brain sections. Data panels **D, E** and **G** represents mean ± SEM (error bars smaller than the data points are not displayed). In **G**, one-way ANOVA (F_(2,15)_s52.79) followed by Dunnett’s multiple comparison tests, *** represents p < 0.0001 compared with basal.

As recently reported^4^, systemic administration of C21 led to a low brain/serum ratios of the compound in mice and a similar result (CSF/serum) was observed in the macaque (**Figs. 2A, B**). These low ratios suggested poor brain penetrance. Unlike CNO^2,5^, C21 was not a substrate for the P-glycoprotein (P-gp) efflux pump (**Extended Fig. 2**). To assess direct brain engagement of DREADDs we injected wild-type (WT) and transgenic mice expressing hM3Dq or hM4Di in dopamine D1 receptor-expressing neurons (D1-hM3Dq or D1-hM4Di) intraperitoneally (IP) with [^3^H]C21 or [^3^H]clozapine and collected brain and organs 30 or 60 min later. [^3^H]C21 showed widespread uptake across various organs and in blood. The highest levels of [^3^H]C21 were observed in kidneys (**Extended Fig. 3**). In brain, [^3^H]C21 accumulated exclusively in ventricles without discernible DREADD engagement (**Figs. 2C-E**). In agreement with the results of the *ex vivo* autoradiography described immediately above, PET imaging revealed that 1 mg/kg C21 (mouse: IP, macaque: IV) led to low (~10%) *in vivo* DREADD occupancy in mice (**Figs. 2F, G**) and no measurable occupancy in the macaque (**Figs. 2H, I**). At 10 mg/kg, C21 achieved ~50% hM4Di occupancy in the macaque (**Figs. 2H, I**) but this dose also produced extensive displacement of [^11^C]clozapine binding at endogenous binding sites and sedative effects, thus no further NHP studies were conducted. C21 did not affect locomotor activity in WT mice at doses up to 1 mg/kg but, like in the macaque, produced sedative effects at 10 mg/kg (**Fig. 2J**). In D1-hM3Dq mice, C21 exhibited a robust decrease of locomotor activity at 0.1 and 1 mg/kg (**Fig. 2K**) and seizure-like behavior at 10 mg/kg. Thus, no further experiments were performed at this dose. In D1-hM4Di mice, C21 produced a small but significant decrease in locomotor activity at 1 mg/kg and sedation similar to that observed in the other groups of mice at 10 mg/kg (**Fig. 2L**). The doses required to induce the same degree of DREADD-specific locomotor inhibition by clozapine were approximately 100-fold lower. DREADD-specific behavioral effects of C21 were also examined in transgenic rats expressing hM3Dq the ventral tegmental area (VTA) (**Fig. 2M**). Rats were injected IP with 1 or 5 mg/kg C21, which led to significant increases in locomotor activity, while clozapine produced comparable locomotor increases at a dose as low as 0.001 mg/kg (**Figs. 2N, O**). DREADD-mediated behavioral effects were observed at low levels of *in vivo* occupancy, likely due to supraphysiological viral-mediated DREADD expression and high intrinsic efficacy of C21 and clozapine.

**Figure 2.**
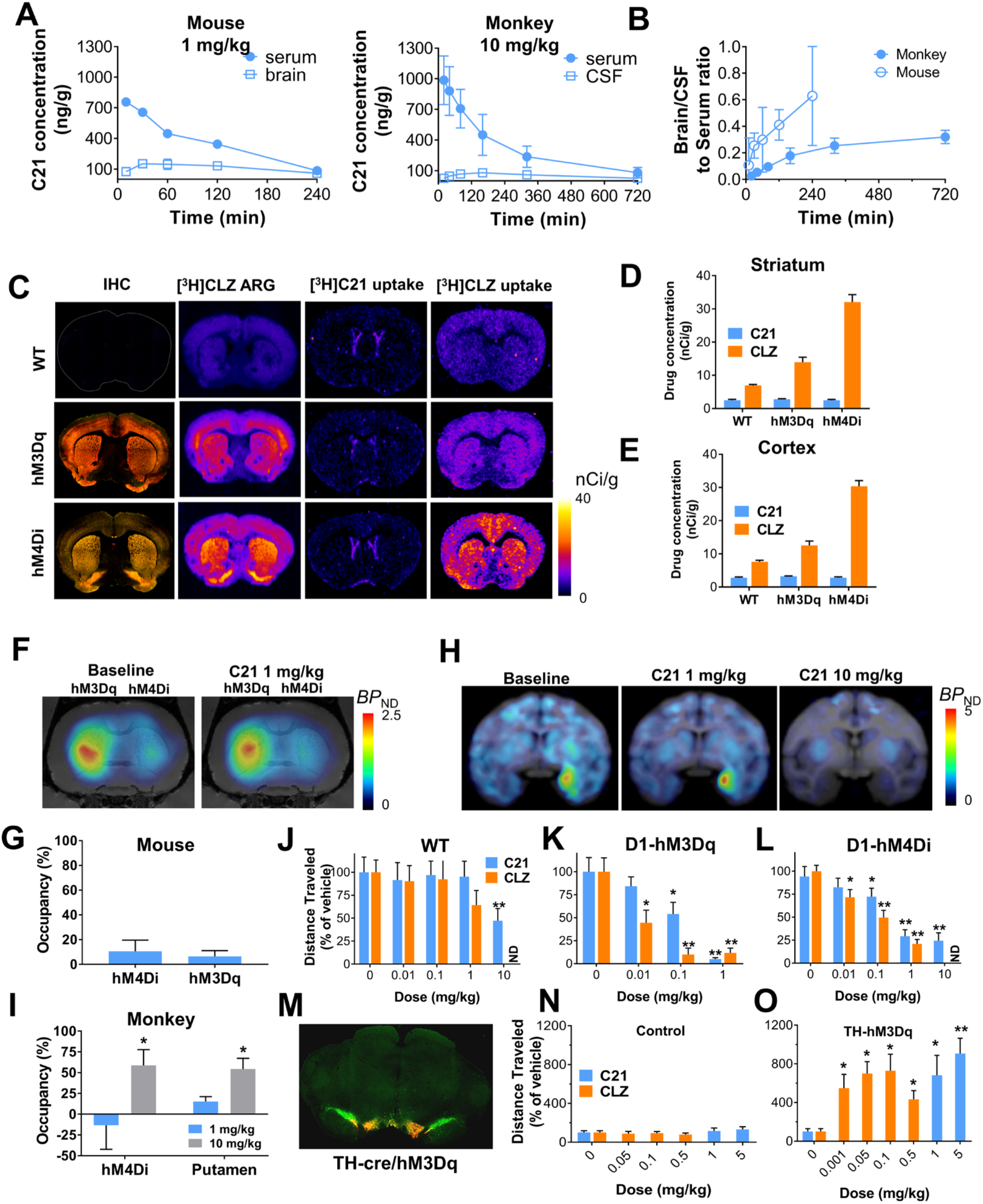
C21 exhibits poor brain penetrance and low *in vivo* DREADD potency in rodents and in nonhuman primates. (**A**) Systemic C21 administration leads to low brain and high plasma concentrations in mice (n = 3 mice) and in macaques (n = 2 macaques) and (**B**) low brain/serum and CSF/serum ratios. (**C-E**) Immunohistochemical (IHC) and autoradiographic (ARG) localization of HA (hemagglutinin)-DREADDs in transgenic mouse brain tissue (non-fused mCitrine (yellow) and fused HA-tag (red)) along with ex vivo [^3^H]C21 and [^3^H]CLZ brain uptake in striatum and cortex showing that [^3^H]C21 does not bind to DREADDs *ex vivo*. Representative images from sections collected from 3 different mice per condition are displayed in **C**, quantification of the uptake in cortex and striatum is displayed in **D** and **E**. (**F, G**) 1 mg/kg C21 (IP) produces an approximate 10% displacement of [^11^C]CLZ in mice (n = 5 mice) expressing AAV-DREADDs in striatum. (**H, I**) Systemic delivery of 1 mg/kg C21 does not displace [^11^C]CLZ binding in macaques (n = 2 macaques) expressing AAV-hM4Di in the amygdala whereas 10 mg/kg displaces [^11^C]CLZ from both hM4Di and non-hM4Di expressing sites. (**J-L**) CLZ exhibits significantly greater *in vivo* potency than C21 in transgenic DREADD mice (n = 8 to 18 mice per condition) and (**M-O**) in transgenic rats (n = 10 rats per condition) expressing hM3Dq in tyrosine-hydroxylase (TH) neurons in ventral tegmental area (VTA). **M** is a representative IHC image (TH green, mCherry red) of the animals used in O. In all cases, data are represented as mean ± SEM. One or two-way repeated measures ANOVA followed by Dunnett’s multiple comparison tests were performed, * p < 0.05 and ** p < 0.01 compared with the respective vehicle.

In a competitive binding screen C21 showed a similar target profile to clozapine at 10 μM, but unlike clozapine, C21 exhibited binding to mu, kappa, and delta opioid receptors (**Extended Fig.4**). C21 tested at 100 nM showed a similar profile to 10 nM of clozapine. However, given that C21 exhibited lower *in vivo* DREADD binding than clozapine, and that higher (~1000-fold) doses needed to drive DREADD-specific behaviors, this *in vitro* selectivity profile for endogenous targets does not appear to predict *in vivo* outcomes very well.

In PET studies using [^18^F]fluorodeoxyglucose (FDG) to measure changes in regional metabolic activity, we injected 1 mg/kg C21 (IP), a dose on the lower end of the range recently suggested for its use^4^. This dose produced significant changes in brain metabolic activity in WT mice (**Extended Fig. 5**), indicating that 1 mg/kg of C21 affects brain function and cannot be used for DREADD-assisted metabolic mapping (DREAMM) studies^6,7^, even though no behavioral effects were detected in our model at this dose. In contrast, an equipotent dose of clozapine (0.1 mg/kg) did not produce any significant changes in brain glucose metabolism (**Extended Fig. 5**).

Collectively, the above findings suggest that C21, like CNO, exhibits suboptimal *in vivo* potency and is not efficient in NHP applications. To search for potent DREADD agonists suitable for *in vivo* biological studies, we profiled two other compounds from the series described in Chen et al.^3^: Compound 13 (C13) and Compound 22 (C22) (**Fig. 3A**). Both C13 and C22 exhibited high DREADD affinity and C13 had 2-fold higher affinity (^hM3Dq^*K*_i_=4.3 nM; ^hM4Di^*K*_i_=4.3 nM) than C22 (^hM3Dq^*K*_i_=11 nM; ^hM4Di^*K*_i_=9 nM) (**Fig. 3B**). Unsurprisingly, given its high *K*_i_, 10 nM C13 was able to displace [^3^H]clozapine from hM3Dq selectively but not from endogenous clozapine binding sites (**Figs. 3C, D**). Next, we synthesized [^3^H]C13 which bound DREADDs expressed in brain tissue sections at very low concentrations and exhibited greater DREADD selectivity than [^3^H]clozapine (particularly in the macaque) (**Figs. 3E-H**). Finally, systemic [^3^H]C13 exhibited high *ex vivo* DREADD engagement in D1-DREADD mice (**Fig. 3I**) indicating high brain permeability and direct DREADD engagement.

**Figure 3.**
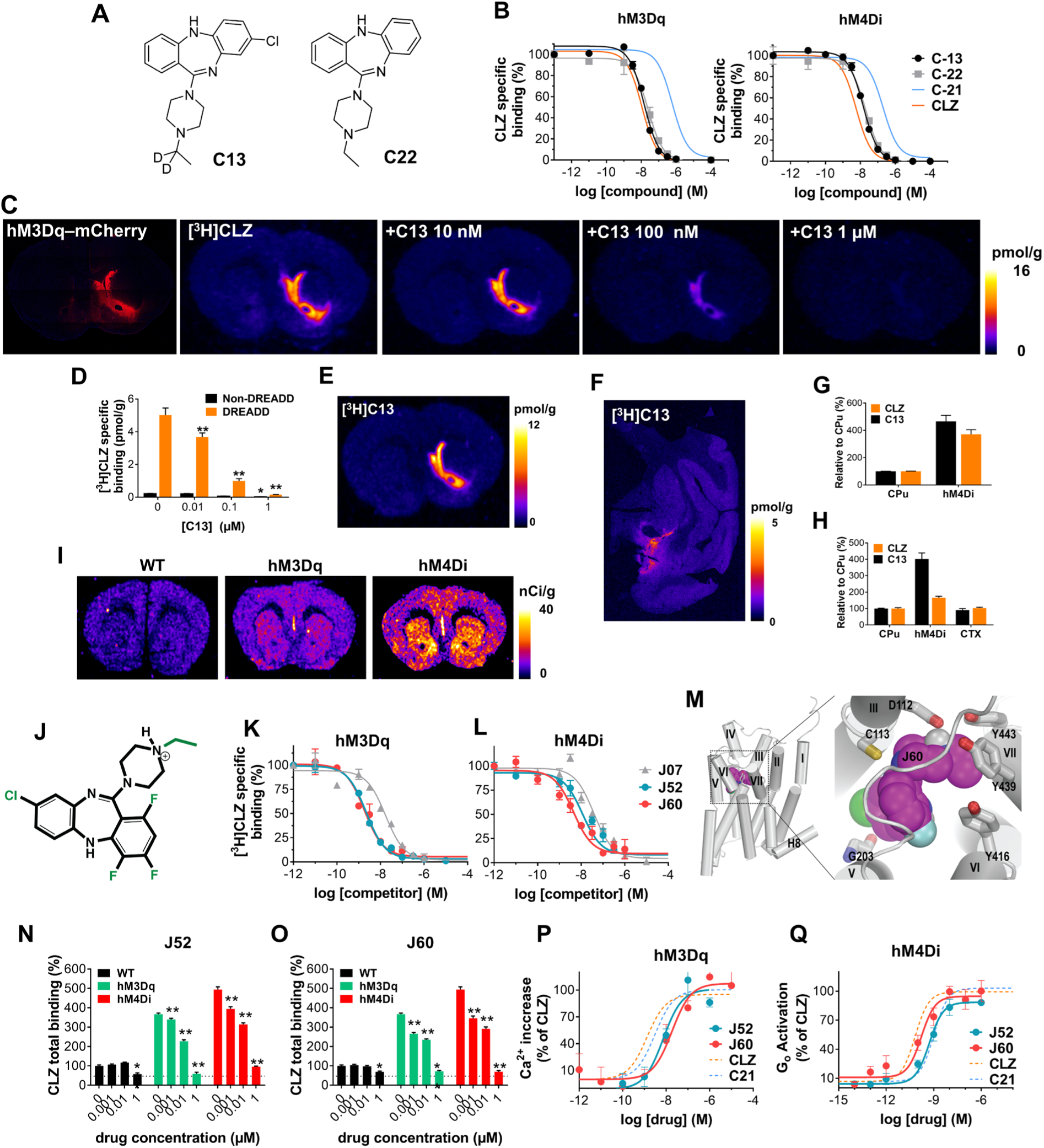
Development and characterization of novel designer ligands displaying high *in vitro* DREADD affinity and potency. (**A**) Compound 13 (C13) and Compound 22 (C22) structures. (**B**) C13 and C22 exhibit comparable DREADD affinity to clozapine (CLZ) with C13 showing ~2-fold greater affinity than C22. CLZ and C21 curves from Figure 1 are overlaid for comparison. (**C, D**) C13 selectively displaces [^3^H]CLZ from AAV-DREADD sites at 10 nM. Representative images of sections collected from 3 different mice are displayed and quantified in **D** as mean ± SEM. Two-way ANOVA followed by Dunnett’s test, * p < 0.05 and ** p < 0.01 compared with the respective vehicle. (**E-H**) [^3^H]C13 binds with greater selectivity than [^3^H]CLZ to DREADDs in mouse and macaque brain tissue expressing AAV-hM3Dq and AAV-hM4Di respectively. (**I**) Intraperitoneal (IP) injection of [^3^H]C13 readily enters the brain to binds to DREADDs. Representative images from 3 mice per condition. (**J-L**) JHU37107 (J07), JHU37152 (J52) and JHU37160 (J60) are high-affinity DREADD ligands. (**M**) Docking and molecular dynamics simulation of J60 in the ligand binding pocket of a hM4Di model. (**N-O**) J60 and J52 selectively displace [^3^H]CLZ at a concentration of 1 and 10 nM from hM3Dq and hM4Di expressed in mouse brain sections (n = 3 mice per condition). (**P-Q**) J60 and J52 activate hM3Dq and hM4Di expressed in HEK293 cells with high potency (experiments performed 3 to 5 times). In all cases, data are represented as mean ± SEM.

The PET radioligand [^11^C]clozapine has been used to image DREADDs *in vivo* in rodents and in NHPs^2,8,9^. However, the short half-life (~20 min) of the ^11^C radionuclide does not permit combined use of chemogenetics with PET at institutions that lack cyclotrons. The ^18^F radionuclide, with a half-life of ~110 min, allows for commercialization, extends the use of PET to chemogenetics applications at institutions that lack cyclotrons, and facilitates imaging at longer time intervals. To make an ^18^F-labeled PET DREADD ligand, we designed and synthesized fluorinated analogs of C13 and C22. The presence of the additional fluorine would make it simple to radiolabel the compound with ^18^F via direct substitution, and by exploring various potential sites for fluorination we would be able to preserve, or perhaps even improve DREADD affinity. We identified three analogs, JHU37107 (^hM3Dq^*K*_i_=10.5 nM; ^hM4Di^*K*_i_=23.5 nM), JHU37152 (^hM3Dq^*K*_i_=1.8 nM; ^hM4Di^*K*_i_=8.7 nM) and JHU37160 (^hM3Dq^*K*_i_=1.9 nM; ^hM4Di^*K*_i_=3.6 nM) displaying the highest *in vitro* DREADD affinity (**Figs. 3J-L**). We then docked the highest-affinity JHU37160 in the ligand binding pocket of an hM4Di model and identified a stable pose of JHU37160 using molecular dynamics pyramidal nitrogen of JHU37160 forms an ionic interaction with Asp112^3.32^, its ethyl group makes favorable hydrophobic-aromatic interactions with Tyr416, Tyr439, and Tyr443, which likely contribute to the ~25-fold improved affinity of JHU37160 compared to C21. The chloride group of JHU37160 forms a halogen bond with the backbone carbonyl oxygen of Gly203 (note Gly203 is one of the two mutations in DREADD). Based on this pose, we predicted that fluorination at the para positions would be well tolerated, which was indeed the case for JHU37152 and JHU37160.

We tested both compounds for *in situ* [^3^H]clozapine displacement in brain tissue from WT and D1-DREADD mice and found that both JHU37152 and JHU37160 exhibited selective [^3^H]clozapine displacement from DREADDs and not from other clozapine binding sites at concentrations up to 10 nM (**Figs. 3N, O**). Additionally, both compounds were potent DREADD agonists with high potency and efficacy in fluorescent and BRET-based assays in HEK-293 cells; JHU37152: (^hM3Dq^*EC_50_*: 5 nM; ^hM4Di^*EC_50_*: 0.5 nM) and JHU37160: (^hM3Dq^*EC_50_*: 18.5 nM; ^hM4Di^*EC_50_*: 0.2 nM) (**Figs. 3P, Q**), whereas no responses were observed in untransfected HEK-293 cells (**Extended Fig. 1**).

In contrast to CNO and C21, mice injected (IP) with a 0.1 mg/kg dose of either JHU37152 or JHU37160 (**Fig. 4A**) showed high brain/serum concentration ratios (~8-fold higher in the brain than serum at 30 min), indicating active sequestration in brain tissue (**Fig. 4B**). Neither JHU37152 nor JHU37160 were P-gp substrates (**Extended Fig. 2**).

**Figure 4.**
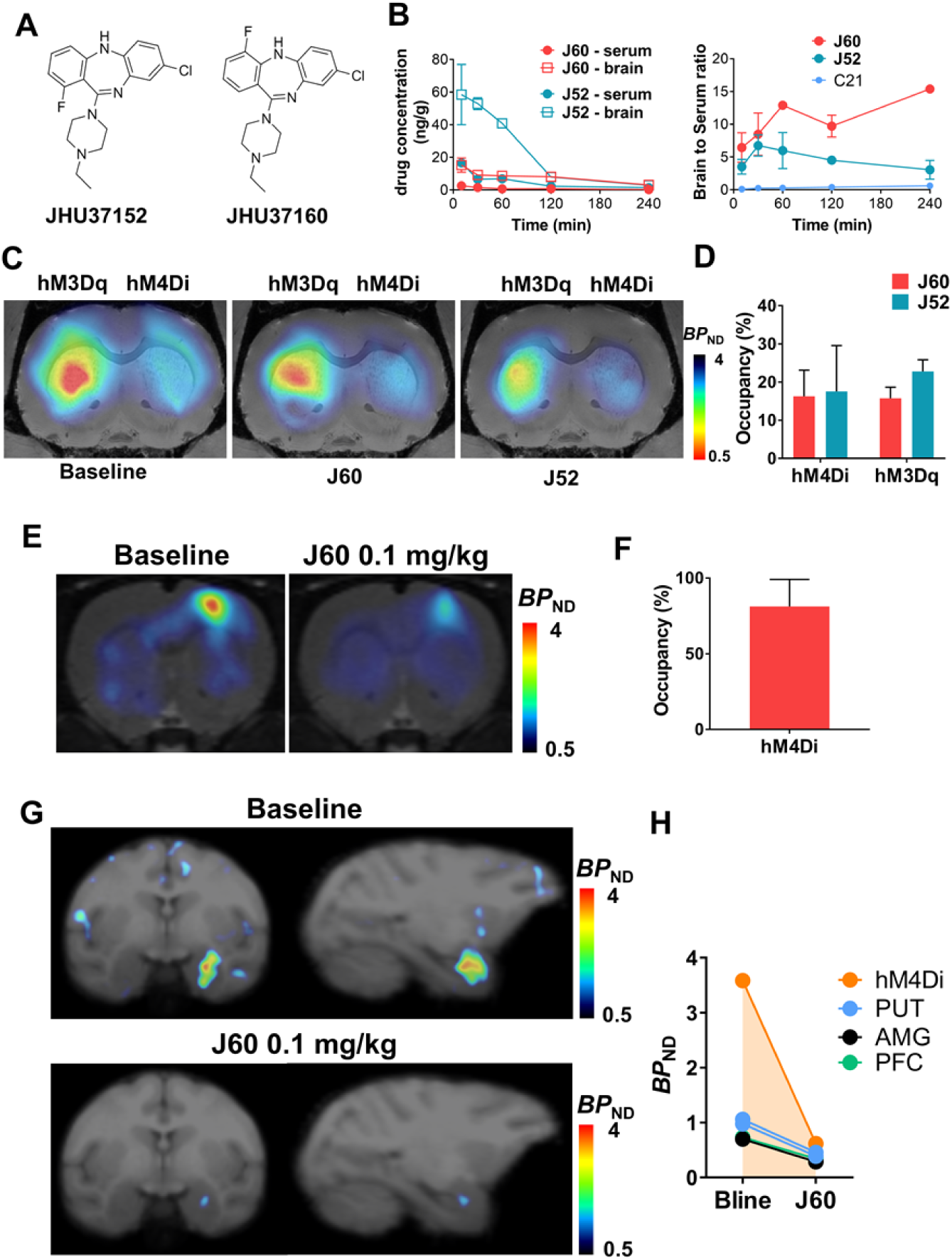
JHU37152 and JHU37160 exhibits high *in vivo* DREADD occupancy. (**A**) Structures of JHU37152 (J52) and JHU37160 (J60). (**B**) Brain/serum concentrations and ratios of J52 and J60 in mice (n = 4 mice per condition) C21 data is same as shown in Figure 2 for comparison purposes. (**C, D**) 0.1 mg/kg of J52 and J60 displace [^11^C]clozapine DREADD binding *in vivo* in AAV-DREADD expressing mice (n = 5 mice). (**E, F**) J60 (0.1 mg/kg, IP) selectively displaces [^11^C]clozapine AAV-DREADD binding *in vivo* in rats (n = 3 rats). (**G, H)** J60 (0.1 mg/kg) displaces [^11^C]clozapine hM4Di binding in the macaque. All data represented are mean ± SEM except in **H** where individual values are displayed.

The CSF concentration of JHU37160 at this same dose in the macaque was below our system detection limit. However, JHU37160 was detected in serum where it showed a similar profile as in the mouse (**Extended Fig. 6**). At this same dose, 0.1 mg/kg, JHU37152 and JHU37160 occupied approximately 15–20% of striatal DREADDs in D1-DREADD mice (**Figs. 4C, D**). In rats, 0.1 mg/kg JHU37160 occupied approximately 80% of cortical hM4Di (**Figs. 4E, F**). In macaques, 0.1 mg/kg JHU37160 (and to a lesser extent, 0.01 mg/kg produced [^11^C]clozapine DREADD displacement at hM4Di expressed in the amygdala (**Figs. 4G, H, and Extended Fig. 6**).

As predicted from the above findings, JHU37152 and JHU37160 were potent *in vivo* DREADD agonists, selectively inhibiting locomotor activity in D1-hM3Dq and D1-hM4Di mice at doses ranging from 0.01 to 1 mg/kg without any significant locomotor effects observed at these doses in WT mice (**Figs. 5A-C**).

**Figure 5.**
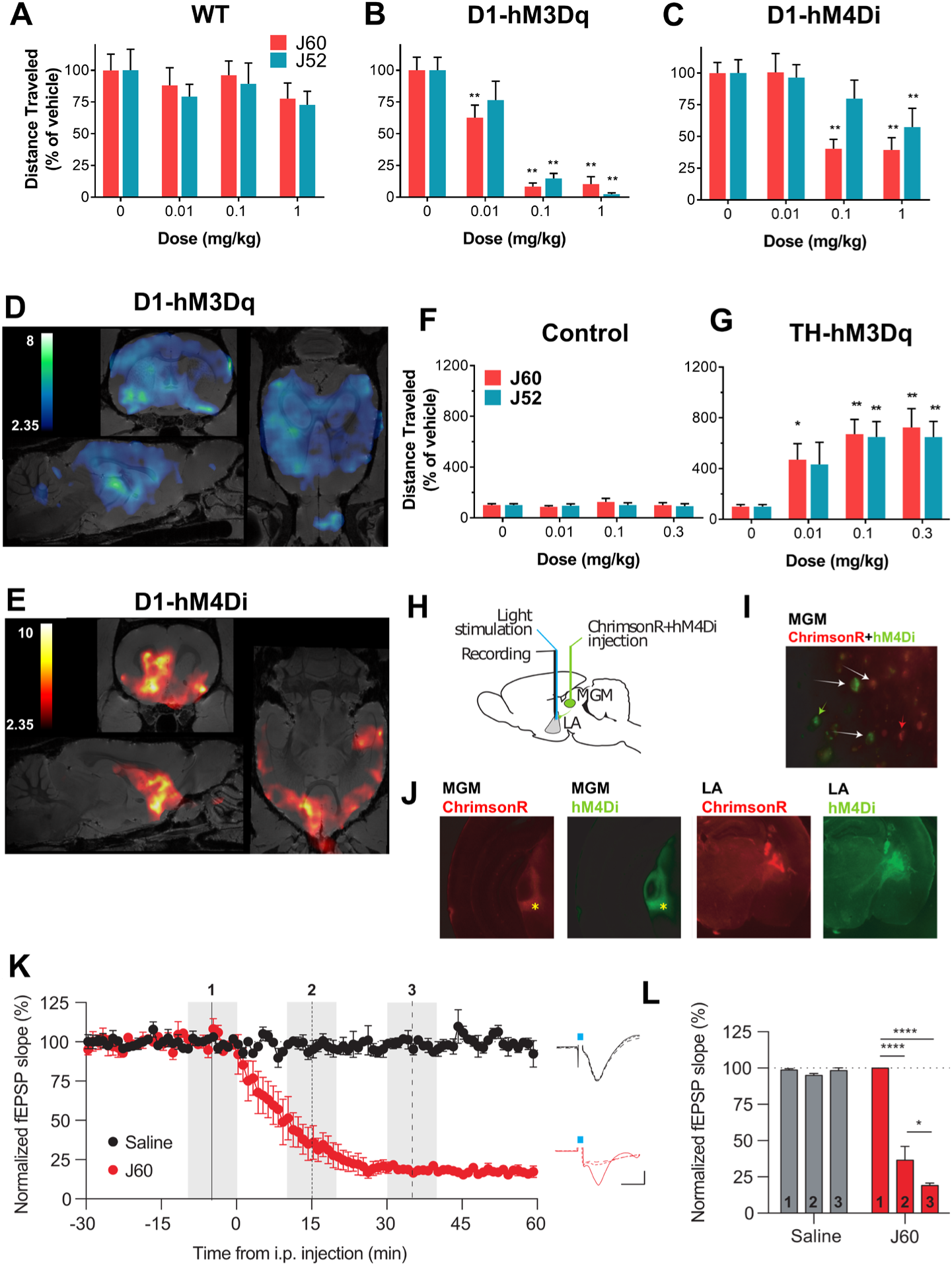
JHU37152 and JHU37160 exhibit high *in vivo* DREADD potency. (**A-C**) J60 and J52 produce potent inhibition of locomotor activity in transgenic D1-DREADD mice but not in wildtype (WT) mice (n = 7 to 19 mice per condition). Two-way repeated measures ANOVA followed by Dunnett’s multiple comparison tests were performed, * p < 0.05 and ** p < 0.01 compared with the respective vehicle. (**D, E**) DREADD-assisted metabolic mapping (DREAMM) using FDG and PET in D1-hM3Dq and D1-hM4Di mice (n = 4 mice per condition) reveals opposing and differential recruitment of whole-brain functional networks. (**F, G**) J52 and J60 produce potent activation of locomotor activity in rats (n = 7 rats per condition) expressing hM3Dq in tyrosine hydroxylase-expressing neurons in the ventral tegmental area. One-way repeated measures ANOVA followed by Dunnett’s multiple comparison tests were performed, * p < 0.05 and ** p < 0.01 compared with the respective vehicle. (**H-J**) Design of *in vivo* electrophysiological experiment and IHC showing hM4Di (green) and ChrimsonR (red) expression in the medial division of the medial geniculate nucleus (MGM) and lateral amygdala (LA). (**K, L**) J60 (0.1 mg/kg) produces rapid and potent hM4Di-driven inhibition of light-evoked neuronal activation. Data are represented as mean ± SEM, * p < 0.05, *** p < 0.001.

Using DREAMM to assess neuronal activity, 0.1 mg/kg (IP) JHU37160 produced metabolic changes in distinct and largely non-overlapping brain networks in D1-hM3Dq (**Fig. 5D**) and D1-hM4Di (**Fig. 5E**) mice and caused no significant brain metabolic changes in WT mice (**Extended Fig. 5**). DREAMM revealed the recruitment of distinct, almost mutually exclusive networks, paralleled by metabolic changes with opposite directionality upon differential modulation of D1 neurons with hM3Dq and hM4Di: decreased metabolism in D1-hM3Dq and increased metabolism in D1-hM4Di mice, effects likely mediated via activation and inhibition of striatal GABAergic D1-expressing neurons respectively.

In a competitive binding screen, JHU37152 and JHU37160 exhibited lower affinity than clozapine at 5-HT receptors (**Extended Fig. 4**). Although the overall target profile of both compounds was similar to clozapine, they did not produce any agonistic effect in functional assays performed in HEK-293 cells lacking DREADDs but expressing endogenous clozapine-binding targets^10^ (**Extended Fig. 1**). As such, they are expected to behave as antagonists at these receptors, competing with endogenous neurotransmitters at these same binding sites. In contrast, JHU37152 and JHU37160 evidenced DREADD activation at lower concentrations than clozapine, indicating that the former compounds are more selective DREADD agonists.

In TH-hM3Dq rats, 0.01–0.3 mg/kg JHU37152 and JHU37160 led to robust, selective increases in hM3Dq-stimulated locomotion (**Figs. 5F, G**). To further characterize the performance of JHU37160 as an *in vivo* DREADD agonist, we performed *in vivo* electrophysiology experiments in which hM4Di was co-expressed with a new light-drivable channelrhodopsin, ChrimsonR (**Extended Fig. 7**), in the terminals of the medial division of the medial geniculate nucleus (MGM) to striatum/lateral amygdala (LA) pathway (**Figs. 5H-J**) in mice. Mice were implanted with optrodes in the LA. A dose of 0.1 mg/kg (IP) JHU37160 elicited rapid inhibition of ChrimsonR-induced terminal activation; 60% inhibition was observed at ~10 min (36 ± 9% of baseline), and maximal inhibition at 30 min after injection (19 ± 2% of baseline) (**Figs. 5K, L**). These effects were hM4Di-dependent; the same injection of JHU37160 had no effect on electrical stimulation evoked responses in animals without DREADD expression (**Extended Fig. 8**).

The high-affinity profiles of JHU37152, JHU37160, and JHU37107 stimulated efforts to develop them into ^18^F-labeled PET imaging probes. The most radiochemically-favorable structure was [^18^F]JHU37107 which we radiolabeled with high yield, molar activity and radiochemical purity (**Fig. 6A**). [^18^F]JHU37107 exhibited robust *in vivo* DREADD binding in D1-DREADD mice and was readily displaceable by 0.1 mg/kg (IP) JHU37160 (**Figs 6B-D**). We also tested [^18^F]JHU37107 in rats with AAV-transduced hM3Dq expression in the unilateral (right) motor cortex (**Fig. 6E**). [^18^F]JHU37107 permitted hM4Di visualization in both the right motor cortex (AAV injection site) as well as at proximal (striatum, contralateral cortex, corpus callosum fibers) and long-range, distal projection sites (thalamus) (**Figs. 6F-H**). [^18^F]JHU37107 exhibited favorable pharmacokinetic properties and metabolite profile in the macaque (Extended Fig. 9) where it was able to directly label hM4Di expressed in the amygdala and at putative axonal projection sites (**Figs. 6I-K**).

**Figure 6.**
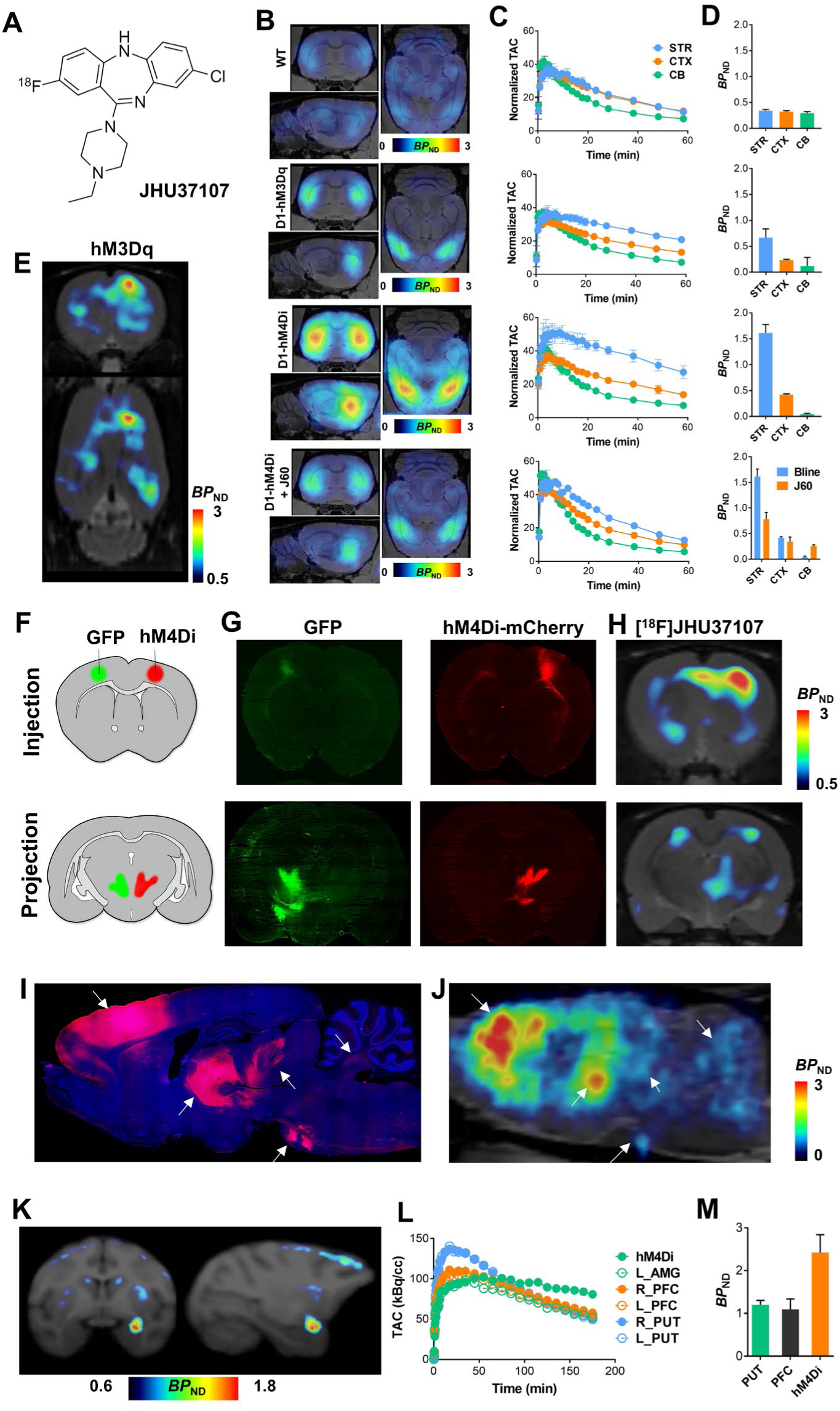
[^18^F]JHU37107 enables non-invasive detection of DREADD in locally-targeted cells and at their long-range projection sites. (**A**) Structure of [^18^F]JHU37107. (**B-D**) [^18^F]JHU37107 selectively binds to DREADDs in the brain of transgenic D1-DREADD mice (n = 3 mice per condition) and is blocked by 0.1 mg/kg of JHU37160. (**E-J**) [^18^F]JHU37107 selectively binds to AAV-DREADDs expressed in the rat cortex and enables non-invasive and longitudinal mapping of both local (injection site) and long-range projections of motor cortex circuitry (ventrolateral thalamus-pontine nuclei shown as main hubs). Representative immunohistochemical images showing GFP (green) or mCherry (DREADD) from a single rat are shown side by side with [^18^F]JHU37107 PET images obtained from the same rat. Arrows used to show corresponding anatomical regions. (**K-M**) [^18^F]JHU37107 binds to hM4Di expressed in the macaque amygdala and at putative projection sites. All data are represented as mean ± SEM except in **L-M** where individual values are displayed.

The human muscarinic receptor-based DREADDs are the most popular chemogenetic technology for basic research and are used by hundreds of laboratories around the world. Although the majority of DREADD use has been in rodents, DREADDs have also been applied to the NHP recently^8,11,12^, a critical step before human translation. Results from prior studies^2,5^, and now from the current study, indicate that the DREADD agonists developed to date, including the most recent agonist, C21, exhibit suboptimal properties and are not appropriate for translational applications. We addressed this here by developing the first DREADD agonists that exhibit high *in vivo* potency and CNS DREADD occupancy that translates from rodents to NHP. We also developed the first, high-affinity ^18^F-labeled DREADD PET ligand with translational potential, which offers the unprecedented ability to visualize DREADD expression at both local and at long-range projection sites, enabling, for the first time, noninvasive and longitudinal visualization of cell type-specific neuroanatomical projections in the living subject.

Chemogenetic technologies, like DREADDs offer the ability to remotely control neuronal activity across distributed brain circuits in a cell-specific manner, without the need for implantable devices^13^, thereby making them especially useful in awake and even unrestrained animals. These characteristics render such tools promising for clinical neurotherapeutics. With the advent of [^18^F]JHU37107 and our new agonists, DREADD affords potential as a neurotheranostic technology, bringing the goals of cell type- and circuit-specific neuromodulation closer. Noninvasive visualization and quantification of DREADDs can be carried out with PET, and with our new ligands, longitudinal neuronal projection mapping and estimation of agonist dose-DREADD occupancy can be assessed. Finally, DREAMM can be used to evaluate, longitudinal, noninvasive assessment of whole-brain, functional circuit activity. If the novel pharmacological tools and approaches we describe here are extended to humans, DREADD-based neurotheranostics would comprise a precision-medicine approach that could be used for informing treatment efficacy parameters of chemogenetic neuromodulation at the individual patient level.

Clinical trials for brain diseases using technologies such as AAV-mediated gene therapy have already been implemented or are currently ongoing and are thus paving the way for safety considerations^14^. Implementation of DREADD-based neurotheranostics in patients would require certain considerations such as safety and efficacy of gene therapy delivery methods, identification of neuron types and circuits to target, and ethical considerations for certain populations. It is important to point out that such considerations are not unique to chemogenetics but are shared by conventional brain-targeted gene therapies. New technologies such as acoustically-targeted chemogenetics^15^ and systemically administered genetically-targeted vectors ^16^ offer further promise for minimizing invasiveness of such practices. Chemogenetics and related technologies are already greatly facilitating identification of disease-relevant neuron types and circuits at the preclinical level, and tools such as those reported here hold great promise for helping translate these groundbreaking studies into clinical strategies. In sum, chemogenetics-based neurotheranostics using the tools we present here offer a novel and promising precision-medicine approach for tackling the most intractable brain disease states underlying the most devastating neurological and psychiatric disorders.

## Methods

### Experimental subjects

Wild-type mice (C57BL/6J) were ordered from Jackson Laboratories and rats (Sprague-Dawley) were ordered from Charles River. Rodents were male and ordered at ~6-weeks of age. Transgenic mice were bred at NIDA breeding facility. Transgenic mice expressing the enzyme cre recombinase under the control of the dopamine D1 receptor promoter (D1-Cre, FK150 line, C57BL/6J congenic, Gensat, RRID: MMRRC_036916-UCD) were crossed with transgenic mice with cre recombinase-inducible expression of hM4Di DREADD (R26-hM4Di/mCitrine, Jackson Laboratory, stock no. 026219) or hM3Dq DREADD (R26-hM3Dq/mCitrine, Jackson Laboratory, stock no. 026220). Three male rhesus monkeys (*Macaca mulatta*) weighed 8 – 12 kg. All experiments and procedures followed NIH guidelines and were approved by each institute’s animal care and use committees.

### Cell culture and transfection

Human embryonic kidney (HEK-293, ATCC) cells were grown in Dulbecco’s modified Eagle’s medium (DMEM; Gibco, ThermoFisher Scientific, Waltham, MA, USA) supplemented with 2 mM L-glutamine, antibiotic/antimycotic (all supplements from Gibco) and 10% heat-inactivated fetal bovine serum (Atlanta Biologicals, Inc. Flowery Branch, GA, USA) and kept in an incubator at 37°C and 5% CO_2_. Cells were routinely tested for myclopasma contamination (MycoAlert^®^ Mycoplasma Detection Kit, Lonza). Cells were seeded on 60 cm^2^ dishes at 4×10^6^ cells/dish 24 h before transfection. The indicated amount of cDNA was transfected into HEK-293 cells using polyethylenimine (PEI; Sigma-Aldrich) in a 1 to 2 DNA:PEI ratio. Cell harvesting for radioligand binding experiments or signaling assays were performed approximately 48 h after transfection.

### Radioligand binding assays

HEK-293 cells were transfected with 5 μg/dish of AAV packaging plasmids encoding for hM3Dq (Addgene #89149), hM4Di (Addgene #89150) or a control vector and harvested 48 hrs after transfection. Cells were suspended in Tris-HCl 50 mM pH 7.4 supplemented with protease inhibitor cocktail (1:100, Sigma-Aldrich, St Louis, MO, USA). The dissected brain tissue was diluted in Tris-HCl 50 mM buffer supplemented with protease inhibitor cocktail (1:1000). HEK-293 cells or brain tissue were disrupted with a Polytron homogenizer (Kinematica, Basel, Switzerland). Homogenates were centrifuged at 48,000*g* (50 min, 4 °C) and washed twice in the same conditions to isolate the membrane fraction. Protein was quantified by the bicinchoninic acid method (Pierce). For competition experiments, membrane suspensions (50 μg of protein/ml) were incubated in 50 mM Tris-HCl (pH 7.4) containing 10 mM MgCl_2_, 2.5 nM of [^3^H]clozapine (83 Ci/mmol, Novandi Chemistry AB, Södertälje, Sweden) and increasing concentrations of the competing drugs during 2hr at RT. Non-specific binding was determined in the presence of 10 μM clozapine. In all cases, free and membrane-bound radioligand were separated by rapid filtration of 500-μl aliquots in a 96-well plate harvester (Brandel, Gaithersburg, MD, USA) and washed with 2 ml of ice-cold Tris-HCl buffer. Microscint-20 scintillation liquid (65 μl/well, PerkinElmer) was added to the filter plates, plates were incubated overnight at RT and radioactivity counts were determined in a MicroBeta2 plate counter (PerkinElmer, Boston, MA, USA) with an efficiency of 41%. One-site competition curves were fitted using Prism 7 (GraphPad Software, La Jolla, CA, USA). K_i_ values were calculated using the Cheng-Prusoff equation.

### *In vitro* functional assays

BRET assays were performed to detect receptor ligand-induced G_α_o1 protein activation. HEK-293 cells were transfected with 5 μg/dish of pAAV plasmids encoding for hM3Dq (Addgene #89149), hM4Di (Addgene #89150) or a control vector together with 0.5 μg Gα-Rluc8, 4.5 μg β1 and 5 μg γ2-mVenus/dish. 48 hours after transfection cells were harvested, washed and resuspended in phosphate-buffered saline (PBS). Approximately 200,000 cells/well were distributed in 96-well plates, and 5 μM Coelenterazine H (substrate for luciferase) was added to each well. Five minutes after addition of Coelenterazine H, ligands were added to each well. The fluorescence of the acceptor was quantified (excitation at 500 nm and emission at 540 nm for 1-second recordings) in a PheraStar FSX plate reader (BMG Labtech) to confirm the constant expression levels across experiments. In parallel, the BRET signal from the same batch of cells was determined as the ratio of the light emitted by mVenus (510–540 nm) over that emitted by RLuc (485 nm). Results were calculated for the BRET change (BRET ratio for the corresponding drug minus BRET ratio in the absence of the drug) 5 minutes after the addition of the ligands.

Intracellular Ca^2+^ concentration was monitored using the fluorescent Ca^2+^ biosensor GCaMP6f. HEK-293 cells were transfected with 7 μg/dish of the cDNA encoding for hM3Dq (Addgene #89149) or hM4Di (Addgene #89150) and 7μg/dish of GCaMP6. 48 h after transfection, cells were harvested, washed, resuspended in Mg^2^+-free Locke’s buffer pH 7.4 (154 mM NaCl, 5.6 mM KCl, 3.6 mM NaHCO3, 2.3 mM CaCl2, and 5 mM HEPES) containing 5.6 mM of glucose and approximately 200,000 cells/well were distributed in black 96-well plates. Increasing concentrations of the indicated compound were added to the cells and fluorescence intensity (excitation at 480 nM, emission at 530 nM) was measured at 18 s-intervals during 250 s using a PHERAstar FSX (BMG Labtech). The net change in intracellular Ca^2+^ concentration was expressed as F-F_0_ where F is the fluorescence at a given concentration of ligand and F_0_ is the average of the baseline values (Fluorescence values of buffer-treated wells).

### Autoradiography

Flash frozen tissue was sectioned (20 μm) on a cryostat (Leica, Germany) and thaw mounted onto ethanol-washed glass slides. Slides were pre-incubated (10 min, RT) in incubation buffer (50 mM Tris-HCl pH 7.4 with 10 mM of MgCl_2_), then slides were incubated (60 min) in incubation buffer containing [^3^H]clozapine (3.5 nM), [^3^H]C21 (10 nM, 41 Ci/mmol, Novandi, Sweden) or [^3^H]C13 (3.5 nM, 13 Ci/mmol, Novandi, Sweden) with or without increasing amounts of the indicated cold ligands (Tocris (clozapine, C21) or custom synthesis). Slides were air dried and placed in a Hypercassette™ (Amersham Biosciences) and covered with a BAS-TR2025 Storage Phosphor Screen (FujiFilm, Japan). The slides were exposed to the screen for 5 to 7 days and imaged using a phosphorimager (Typhoon FLA 7000; GE Healthcare).

### [^35^S]GTP*γ*S autoradiography

Flash frozen tissue was sectioned (10 μm) on a cryostat (Leica) and thaw mounted on ethanol cleaned glass slides. Sections were encircled with a hydrophobic membrane using a PAP pen (Sigma-Aldrich). Pre-incubation buffer was pipetted onto each slide and allowed to incubate for 20 min (500 mM Tris-HCl, 100 mM EDTA, 500 mM MgCl_2_ and 4 M NaCl). The pre-incubation buffer was removed via aspiration and each slide was loaded with GDP in the presence of DPCPX and allowed to incubate of 60 min (Pre-incubation buffer, 48 mM GDP, 24 mM DPCPX, Millipore water). GDP buffer was removed via aspiration and [^35^S]GTPγS cocktail (GDP buffer, 20 mM DTT, 48 mM GDP, 24 mM DPCPX, 300 nM [^35^S]GTPγS) with agonists of interest (C21 10 nM, clozapine 10 nM), without agonists (basal condition), or with a saturated concentration of non-radioactive GTP (for non-specific binding) was pipetted onto each slide and allowed to incubate for 90 min. The [^35^S]GTPγS cocktail was removed via aspiration and slides were washed in ice cold washing buffer (50 mM Tris-HCl, 5 mM MgCl_2_, pH 7.4) for 5 min (2×) followed by a 30 second dip in ice-cold deionized water. Hydrophobic membrane was removed with a cotton swab and xylene and slides were placed into a Hypercassette™ covered by a BAS-SR2040 phosphor screen (FujiFilm; GE Healthcare). The slides were exposed to the phosphor screen for 3 to 5 days and imaged using a phosphor imager (Typhoon FLA 7000; GE Healthcare).

### Binding and enzyme target profile screen

These experiments were performed by an outside vendor (Eurofins, France). Briefly, membrane homogenates from stable cell lines expressing each receptor/enzyme were incubated with the respective radioligand in the absence or presence of clozapine or C21 or reference control compounds in a buffer. In each experiment, the respective reference compound was tested concurrently with the test compound to assess the assay reliability. Nonspecific binding was determined in the presence of a specific agonist or antagonist at the target. Following incubation, the samples were filtered rapidly under vacuum through glass fiber filters presoaked in a buffer and rinsed several times with an ice-cold buffer using a 48-sample or 96-sample cell harvester. The filters were counted for radioactivity in a scintillation counter using a scintillation cocktail.

### P-glycoprotein (P-gp) substrate assay

These experiments were performed by an outside vendor (Eurofins, France). C21 was tested in P-gp substrate assessment assays at 10 μM. The A to B and B to A permeability was measured in Caco-2 cells in the presence and absence of verapamil, a P-gp inhibitor. Efflux ratios (*E*) were calculated based on the apparent B-A and A-B permeability with and without verapamil. In each experiment, the respective reference compound was tested concurrently with the test compound to assess the assay reliability. Fluorescein was used as the cell monolayer integrity marker. Fluorescein permeability assessment (in the A-B direction at pH 7.4 on both sides) was performed after the permeability assay for the test compound. The cell monolayer that had a fluorescein permeability of less than 1.5 × 10^−6^ cm/s for Caco-2 was considered intact, and the permeability result of the test compound from intact cell monolayer was reported.

### Bioanalytical methods

Monkey blood and CSF samples were collected from totally implanted subcutaneous access ports (Access Technologies, Virginia), connected to catheters indwelling in the femoral artery or intrathecal space of the spinal column, respectively. Rodent blood samples and brains were collected immediately following sacrifice at the indicated time points after intraperitoneal injection (10 ml/kg) in buffered saline. CSF was immediately frozen on dry ice and stored at −80 °C. Blood samples were allowed to coagulate for 15 minutes and then centrifuged at 4°C for 15 minutes. Serum was collected from the supernatant and stored at a minimum of −30°C until extraction. To 25 μl of serum, 5 μl of internal standard and 110 μl of methanol were added. Samples were centrifuged for 10 minutes at 16,200 × *g* at 4°*C* and the supernatant was transferred to the autosampler vial for analysis. Brains were cut in half and weighed prior to sample preparation. Half brains were homogenized in 490 μl of 85% ethanol: 15% water containing 0.1% formic acid and 5 μl of internal standard using a polytron homogenizer and centrifuged for 10 minutes at 16,200 × *g* at 4°C. 300 μl of supernatant was dried under a stream of nitrogen and resuspended in 150 μl methanol. The resuspended solution was then centrifuged and 100 μl of supernatant was transferred to the autosampler vial for analysis.

Data was acquired using a Nexera XR HPLC (Shimadzu) coupled with a QTRAP 6500 (SCIEX) and was analyzed with Analyst 1.6 (SCIEX). The positive ion mode data was obtained using multiple reaction monitoring (MRM). The instrumental source setting for curtain gas, ion spray voltage, temperature, ion source gas 1, ion source gas 2 were 30 psi, 5500 V, 500 C, 650 psi, and 5560 psi, respectively. The collision activated dissociation was set to medium and the entrance potential was 10 V. C21 was monitored using the MRM ion transition (278.80→166.10) with declustering potentials (DP)=90V, collision cell exit potentials (CXP)=10V and collision energies (CE)=50V. JHU37160 and JHU37152 were monitored using the MRM ion transitions (359.10→288.10) with DP=70V, CXP=8V and CE =28V. Clozapine was monitored using the MRM ion transitions (327.30→270.10) with DP=100V; 80V, CXP=11V and CE=40 V.

Separation of the C21, JHU37160, JHU37152 and clozapine was accomplished using a C18 Security guard cartridge (4.6 × 4 mm) and an Eclipse XDB-C18 column (4.6 × 250 mm, 5 μm, Agilent) at 35°C. Mobile phase A consisted of water containing 0.1% formic acid and mobile phase B was methanol containing 0.1% formic acid. The following linear gradient was run for 21.0 min at a flow rate of 0.4 ml/min: 0–2.00 min 20% B, 7.0 min 80% B, 12 min 90% B, 18.0 min 90% B, 18.1 min 20% B. 12-point Calibration curves were prepared in standard solution by a 0.5 serial dilution of standards from 0.92 μg/ml for C21; 1 μg/ml for JHU37160; and 0.2 μg/ml for JHU37152 and 0.4 μg/ml for clozapine. The injection volume per sample was 10 μl. Samples were kept at 4 °C in the autosampler tray prior to injection.

The data was measured using standard curves and quality controls, but it was not validated to ICH guidelines. The concentrations of C21, JHU37160 and JHU37152 was measured using area ratios calculated with the internal standard clozapine (5 μl of 100 μg/ml) and the concentrations of clozapine was measured using area ratios calculated with JHU37160 as the internal standard (5 μl of 50 μg/ml). Quality control standards (low, middle and high) were prepared by adding the spiking standard to solution to 25 μl of serum and/or a half-brain and relative values are reported.

### Adeno-associated virus (AAV) injections in rodents

Animals were anesthetized with isoflurane or a mix of ketamine/xylazine and prepped on a stereotaxic apparatus (Kopf, Germany). The following AAVs expressing hM4Di or hM3Dq fused to mCherry (or EGFP as a control) under the control of a hSyn promoter were used when indicated: hSyn-hM4D(Gi)-mCherry (Addgene: 50475-AAV8), hSyn-hM3D(Gq)-mCherry (Addgene: 50474-AAV8), hSyn-DIO-hM3D(Gq)-mCherry (Addgene: 44361-AAV8) and hSyn-EGFP (Addgene: 50465-AAV8). Based on corresponding mouse and rat brain atlases (Paxinos and Watson), the following coordinates were used to target: the dorsal striatum: Mouse – AP=1.00, ML=±1.50, DV=-3.55; Rat – AP=1.70, ML=±2.50, DV=-5.50, rat cortex: AP=1.70, ML=±2.50, DV=-3.50, Rat VTA: AP=-5.5, ML=±0.8, DV=-8.2. In all cases 1 μl/side was injected and all injections were performed using a Hamilton Neuros 33G syringes at a flow rate of 50 nl/min, except for the injections targeting the rat VTA, that were injected with a picospritzer over 90 seconds.

### Lentivirus injections in monkeys

Surgical procedures were performed in a veterinary operating facility under aseptic conditions. Vital signs were monitored throughout the procedure. A pre-operative T1-weighted MRI for each monkey was used to determine the stereotaxic coordinates for the sites of the lentivirus injection in the right amygdala. The skull region above the target site was exposed by retracting the skin, fascia, and muscle in anatomical layers. A small region of cranial tissue was then removed (~1.5 cm diameter) to access the dura mater, into which incisions were made to provide access for the infusion apparatus.

Lentivirus expressing an hM_4_Di-CFP fusion protein under an hSyn promoter ^11^ with a titer of >10^9^ infectious particles was loaded into a 100 μL glass syringe (Hamilton Co., MA). The 31-gauge needle of the syringe was sheathed with a silica capillary (450 μm OD) to create a step 1 mm from the base of the aperture. The syringe was mounted in a Nanomite pump (Harvard Apparatus, Cambridge, MA). The needle was lowered through the incision in the dura mater to each of the pre-calculated target sites and 10 or 20 μL was infused at a rate of 1 μL/min. Each monkey received a total of between 10 – 12 injections (for a total injection volume of 120 to 240 μL). Post-infusion, the needle was left in situ for 10 minutes after each injection to allow pressure from the infusate to dissipate. The needle was then slowly removed. At the completion of the injection series the soft tissues were sutured together in anatomical layers.

### Immunohistochemistry

Animals were anesthetized with a ketamine and xylazine mixture and transcardially perfused with PBS followed by 4% paraformaldehyde (PFA). The brains were post-fixed in 4% PFA (overnight, 4 °C) and then placed in 30% sucrose for 3–4 days. The brains were frozen and sectioned on a cryostat (40 μm) and collected in PBS with 0.1% Tween-20 (washing buffer). Slices were blocked with BSA 3% in washing buffer (blocking buffer, 2h RT), and incubated the primary antibody mixture: chicken anti-GFP (1:400, ab13970, Abcam Inc.) and rabbit anti-HA (1:400, C29F4, Cell Signaling Technologies) overnight at 4 °C. Sections were washed in washing buffer (3 × 10 min, RT) and incubated with the secondary antibody mix: Alexa488-conjugated goat anti-rabbit (1:200, A11034, Invitrogen) and Alexa546-conjugated goat anti-chicken (1:400, A11040, Invitrogen) and To-Pro3 iodide (Invitrogen) as a nuclear counterstain. After washing (3 × 10 min in washing buffer and 1 × 5 min in PBS) slices were mounted on glass slides. Alternatively, flash frozen tissue was sectioned (20 μm) on a cryostat (Leica) and mounted on ethanol-soaked glass slides. Sections were fixed with paraformaldehyde (4%, 10 min at RT), permeabilized with PBS with TritonX-100 (0.1%, washing buffer), blocked with bovine serum albumin 5% in washing buffer (2 hours, RT), and then incubated overnight at 4°C with the primary antibodies mixture: rabbit anti-mCherry (1:500, ab167453, Abcam Inc.) and chicken anti-GFP (1:2000, ab13970, Abcam Inc.). Sections were then washed again and incubated with Alexa 647 goat anti-rabbit (1:200, A21245, Invitrogen) and Alexa 488 goat anti-chicken (1:400, A11039, Invitrogen) and washed again. In both cases, sections were coverslipped using mounting medium (ProLong Diamond antifade mountant, Invitrogen), and images were acquired either using a confocal microscope (Examiner Z1, Zeiss, Germany) with a laser scanning module (LSM-710, Zeiss, Germany) or a Leica Zoom.V16 stereo microscope (Leica, Germany).

### Synthesis of [^11^C]clozapine and [^18^F]JHU37107

[^11^C]clozapine was synthesized using the methods developed by Bender et al.^17^ with minor modifications. Briefly, 1 mg of *N*-desmethylclozapine was dissolved in 200 μL of acetonitrile. [^11^C]Methyl triflate was bubbled into the solution until the radioactivity reached a plateau. The reaction was kept at room temperature for 2 min. The solution was then diluted with 200 μL of 40:60 (v:v) acetonitrile:water (0.1% ammonium hydroxide) and injected onto semi-preparative HPLC. The column (Waters XBridge C18 10 mm × 150 mm) was eluted with 40:60 (v:v) acetonitrile:water (0.1% ammonium hydroxide) at a flow rate of 10 mL/min. The radioactive peak corresponding to [^11^C]clozapine (t_R_ = 6.1 min) was collected in a reservoir containing 50 mL of water and 250 mg of L-ascorbic acid. The diluted product was loaded onto a solid phase extraction cartridge (Waters Oasis HLB plus light) and rinsed with 3.0 mL of water. The product was eluted with 400 μL of ethanol into a sterile, pyrogen-free bottle and diluted with 4.0 mL of saline. A 10 μL aliquot of the final product was injected onto an analytical HPLC column (Waters XBridge C_18_ 4.6 mm × 100 mm) and eluted with 35:65 (v:v) acetonitrile:water (0.1% ammonium hydroxide) at a flow rate of 2 mL/min. The radioactive peak corresponding to [^11^C]clozapine (t_R_ = 9.2 min) coeluted with a standard sample. The semi-preparative HPLC eluant used was 25:75 (v:v) acetonitrile:water (0.1% ammonium hydroxide), and no ascorbic acid was added to the reservoir. The molar activity for both [^11^C]clozapine ranged from 351 to 483 GBq/μmol (9,482–13,041 mCi/μmol) at end of synthesis.

[^18^F]JHU37107 was prepared via the no-carrier-added ^18^F-fluorination using an FDG Nuclear Interface module (Muenster, Germany). Briefly, 4 mg precursor (8-chloro-11-(4-ethylpiperazin-1-yl)-5H-dibenzo[b,e][1,4]diazepin-2-ol) and 8 mg tris(acetonitrile)cyclopentadienylruthenium(II)hexafluorophosphate (STREM, Boston, Massachusetts) were dissolved in ethanol (0.3 mL), the solution was heated at 85°C for 25 min and evaporated to dryness under a stream of argon gas. The residue was dissolved in DMSO (0.5 mL) and acetonitrile (0.5 mL) and the solution was added to a dry complex of [^18^F]fluoride and 9 mg 1,3-bis(2,6-di-i-propylphenyl)-2-chloroimidazolium chloride (STREM). The reaction mixture was heated at 130°C for 30 min, diluted with mixture of 0.5 mL acetonitrile, 0.5 mL water and 0.03 mL TFA and injected onto a semi-preparative HPLC column (Luna C18, 10 micron, 10mm × 250 mm) and eluted with 23:77 (v:v) acetonitrile:water (0.1% trifluoroacetic acid) at a flow rate of 10 mL/min. The radioactive peak corresponding to [^18^F]JHU37107 (t_R_ = 9.9 min) was collected in a reservoir containing 50 mL of water and 3 mL aq 8.4% NaHCO_3_ solution. The diluted product was loaded onto a solid phase extraction cartridge (Waters Oasis HLB plus light) and rinsed with 10 mL sterile saline. The product was eluted with 1000 μL of ethanol through a sterile 0.2 μm filter into a sterile, pyrogen-free vial and 10 mL saline was added through the same filter. The final product [^18^F]JHU37107 was then analyzed by analytical HPLC (Luna C18, 10 micron, 4.6mm x 250 mm; mobile phase 23:77 (v:v) acetonitrile:water (0.1% trifluoroacetic acid), flow rate of 3 mL/min; t_R_ = 6.4 min) using a UV detector at 254 nm to determine the radiochemical purity (>95%) and specific radioactivity (152 – 188 GBq/μmol (4,100 – 5,070 mCi/μmol)) at the time synthesis ended.

### [^11^C]clozapine and [^18^F]JHU37107 imaging using positron emission tomography (PET)

Mice and rats were anesthetized with isoflurane and placed in a prone position on the scanner bed of an ARGUS small animal PET/CT (Sedecal, Spain) or a nanoScan PET/CT (Mediso, USA) injected intravenously (~100–200 μL) with [^11^C]clozapine (~700 μCi) or [^18^F]JHU37107 (~350 μCi) and dynamic scanning commenced. When indicated, animals were pretreated with vehicle or the indicated drug 10 min before the injection of the PET radiotracer. Total acquisition time was 60 min.

All macaque studies were acquired dynamically on the Focus 220 PET scanner (Siemens Medical Solutions, Knoxville, TN). The Focus 220 is a dedicated pre-clinical scanner with a transaxial FOV of 19 cm and an axial FOV of 7.5 cm. Image resolution is < 2 mm within the central 5 cm FOV.

After initial evaluation the monkey was sedated with ketamine (10 mg/kg) followed by ketoprofen (as an analgesic) and glycopyrrolate (for salvia reduction), all weight dependent IM injections. The monkey would then be placed in the supine position, intubated with a tracheal tube. Anesthesia was maintained by 1–3.5% isoflurane and oxygen, the monkey’s head was positioned and immobilized for optimal positioning of the brain and moved into the scanner. A 10-minute transmission scan using a Co-57-point source for attenuation correction was performed. One iv was inserted, if possible, in the right arm and one in the right leg for injection of tracer and blocking agent. The monkey was always monitored while anesthetized. HR, BP, O_2_ saturation, RR, 3 lead ECG, rectal temperature was documented every 15 minutes.

Arterial blood sampling was acquired throughout the study using an indwelling femoral port. The first two minutes samples were collected every 15 seconds then at 3, 5 10,30, 60, 90, and 120 minutes post tracer injection. All scans were acquired for 120 minutes using list mode acquisition.

Scan data were histogrammed into 33 frames (6 ×30 seconds, 3 ×1 minute, 2×2 minutes and 22 ×5 minutes). Reconstruction was performed by Filtered Back Projection with scatter correction. After completion of the last study of the day, isoflurane was cut off. The monkey was gradually awakened, moved to the housing facility and fully recovered.

In all cases, the PET data were reconstructed and corrected for dead-time and radioactive decay. All qualitative and quantitative assessments of PET images were performed using the PMOD software environment (PMOD Technologies, Zurich Switzerland). Binding potential *BP*_ND_ (a relative measure of specific binding) was calculated using a reference tissue model using the cerebellum as a reference tissue in rodents. In macaques, the kinetic data were fitted to a two-tissue compartment model and the concentration of parent in plasma was used as an input function, then the volume of distribution ratios compared to cerebellum were calculated to establish *BP*_ND_. In all cases, the dynamic PET images were coregistered to MRI templates and time-activity curves were generated using predefined volumes of interest (macaques) or manually drawn in rodents and the described analyses were performed. Receptor occupancy was calculated using the formula: 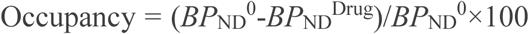, where *BP*_ND_^0^ is the binding potential of the baseline condition and *BP*_ND_^Drug^ the binding potential when the animals were pretreated with the drug. In an independent manner, *BP*_ND_ parametric maps were generated by pixel-based kinetic modeling using a multilinear reference tissue model ^18^ using the cerebellum as a reference region and the start time (t*) was set to 16 min.

The arterial input function for the radiotracers injected in rhesus monkeys was determined using the general methods as previously described ^19^. The radioligand concentrations in the arterial plasma were corrected by the unchanged parent fraction.

Heparinized blood samples (0.5 mL each) were drawn at 15-s intervals until 2 minutes, and at 3, 5, 10, and 30 min followed by 3-mL samples at 60, 90, 120 min ([^11^C]clozapine) and 150, 180 min ([^18^F]JHU37107). Blood samples were immediately sampled for gamma counting, and plasma harvested by centrifugation for gamma counting and radio-HPLC analysis. The unchanged plasma parent fractions were determined by radio-HPLC on an X-terra^®^ *C*18 column (10 *μ*m, 7.8 mm × 300 mm, Waters Corp., Milford, MA), and eluted with MeOH:H_2_O:Et_3_N (80:20:0.1; by volume) at an isocratic flow rate of 4.0 mL/min. Eluates were monitored with an in-line flow-through NaI_(_*_Tl_*_)_ scintillation detector (Bioscan). Data were stored and analyzed on a PC using the software “Bio-ChromeLite”. The collection of data for each radiochromatogram was decay corrected according to its respective HPLC injection time. Plasma free fraction was determined according to our previous methods ^20^.

### DREADD-assisted metabolic mapping (DREAMM)

Mice (D1-hM3Dq, D1-hM4Di or WT littermates, see above) were habituated to experimenter handling and fasted 16 hours before the experiment. On the day of the experiment, mice received an IP injection of vehicle (1 ml/kg) and were placed back into their home cages. Ten minutes later, mice were injected (IP) with 11 MBq of 2-deoxy-2-[^18^F]fluoro-D-glucose (FDG, Cardinal Health) and placed back into their home cages. After 30 minutes, mice were anesthetized with 1.5% isoflurane, placed on a custom-made bed of a nanoScan small animal PET/CT scanner (Mediso Medical Imaging Systems) and scanned for 20 min on a static acquisition protocol, followed by a CT scan. One week later the animals were fasted overnight, the next day received an IP injection of C21 (1 mg/kg), clozapine (0.1 mg/kg) or JHU37160 (0.1 mg/kg) and the FDG-PET procedure was conducted as described above. In all cases, the PET data were reconstructed and co-registered to an MRI template as described above. Voxel-based repeated measures with Student’s t-test comparing baseline to drug were performed, and the resulting parametric images were filtered for statistically significant (p < 0.05) clusters larger than 100 contiguous voxels. All statistical parametric mapping analyses were performed using Matlab R2016 (Mathworks) and SPM12 (University College London).

### *Ex-vivo* biodistribution and brain uptake of [^3^H]clozapine, [^3^H]C21 and [^3^H]C13

Mice were injected (IP) with [^3^H]clozapine, [^3^H]C21 or [^3^H]C13 (2 μCi/g), euthanized 30 min later and brain, blood, and tissues were collected for radiometric analyses. The brains were flash frozen in isopentane (Sigma-Aldrich) and stored at −80°C until use. The blood was centrifuged (13,000 rpm, 10 min at RT) and serum was collected. The tissues were solubilized with Solvable™ (PerkinElmer) and bleached with hydrogen peroxide (Sigma-Aldrich). Serum and tissue samples were dissolved in scintillation cocktail and radioactivity counts were determined in a Beckman LS 60000TA scintillation counter (BeckmanCoulter, Indianapolis, IN, USA).

### Locomotor activity assessment in mice

Transgenic male and female mice (see above) (20–30g) expressing hM3Dq or hM4Di and mCitrine reporters (or controls) were tested for locomotor activity. Mice were injected (IP) with the indicated dose of clozapine, C21, JHU37152 or JHU37160 or vehicle (buffered saline). Ten minutes after injection, animals were placed in an open field arena (Opto-varimex ATM3, Columbus Instruments) and their locomotor activity was tracked during 60 min as infra-red beam crossings and traveled distance was converted to cm. Animals were repeatedly tested on consecutive sessions (after an initial habituation session with no drug treatment) in a counterbalanced design.

### Locomotor activity assessment in rats

Tyrosine hydroxylase (TH):Cre rats (n=9; founder, K. Deisseroth lab) or WT littermates (n=9) were bilaterally injected with AAV2 hSyn-DIO-hM3Dq-mCherry (1ul/hemisphere, Addgene), and allowed to recover for 2+ weeks. Following handling, they were habituated to a Med Associates locomotor testing box (43 × 43 × 30.5 cm) for 2h on 2 days, then repeatedly tested after separate day doses of DREADD agonists and vehicle. Cohort one was administered C21 (1 or 5mg/kg) or clozapine (0.001 to 0.5 mg/kg), and vehicle, in counterbalanced order on separate days, with each test separated by at least 48h. The 2h locomotor testing session commenced 30min after each IP injection on each day. Vehicle was administered on 2 separate days, and average locomotion was used for comparison with DREADD agonists. Cohort 2 underwent the same procedures, but with a 4h testing session, and the following IP injections: vehicle (x2), JHU37152 (0.01 to 0.3 mg/kg), JHU37160 (0.01 to 0.3 mg/kg). All drugs were dissolved in 5% DMSO in saline, and injected 1ml/kg. Following all tests, rats were transcardially perfused, brains sectioned coronally at 40um, and VTA-specific expression confirmed using endogenous mCherry reporter expression, which co-localized nearly exclusively with TH immunoreactivity as previously reported ^21^.

### ChrimsonR generation and characterization

All expression and cloning were similar to previous described ^22^. In brief, pcDNA3.1(+) / Hygro vector (Life Technologies, Carlsbad, CA, USA) was used for expression of Chrimson, oC-Chrimson and oC-Chrimson-ts in HEK293A cells (Life Technologies, Carlsbad, CA, USA). Channelrhodopsin inserts (with and without modification) were cloned into the BamHI and XhoI site and the fluorescent protein (tdTomato or citrine) was cloned in-frame and a 3’ stop codon between the XhoI and XbaI site. For neuron expression, the channelrhodopsin-FP inserts were placed in an AAV2 vector between BamHI and HindIII sites (Addgene #50954). Transfection in HEK293 cells were achieved with Fugene (Roche, Basel, Switzerland) and electroporation (Lonza, Gaithersburg, MD, USA) was used for expression in neuronal culture. HEK293 cell experiments were performed 2 days post-transfection whereas neuronal experiments were performed > 10 days post-transfection. Cell culture conditions were as described previously ^22^.

For visualizing the channelrhodopsin-FP expression, images were taken on a Zeiss Axiovert 200M microscope (Zeiss, Jena, Germany) with Slidebook software (3i, Denver, CO, USA) using a Cascade II 1024 EMCCD camera (Photometrics, Tucson, AZ, USA). Images were taken with a 40x oil objective with NA of 1.2. Citrine images were acquired with 495/10 excitation filter and 535/25 nm filters and tdTomato images were acquired with 580/20 nm excitation and 653/95 nm emission filters (Semrock, Rochester, NY, USA). For analyzing membrane / cytosolic fluorescence, a line profile was drawn across the cell in ImageJ or Fiji, the fluorescence intensity of the 2 membrane regions and cytosolic portion (including nucleus) were measured from profile. After background subtraction, the mean membrane and mean cytosolic fluorescence were calculated and the ratio was calculated.

Electrophysiological recording was performed with identical condition and solutions as previously described ^22^. Due to the desensitization of the Chrimson response to high intensity of light to some wavelengths, a 1 second 410 nm conditioning light was used to illuminate the recorded cell 10 s prior to testing with indicated wavelength of light.

### *In vivo* electrophysiology

Mice were purchased from Janvier Labs and stored in grouped cages (max. 4 per cage) with ad libitum access to food and water, and in a 12h light/dark cycle (lights on at 7:00 am). All the experiments were performed in accordance to the national Danish law for the use of laboratory animals and approved by local authorities.

For AAV injection, mice were anaesthetized with Isoflurane and placed in a stereotaxic holder (Narishige, Japan). A trephine hole was drilled above the MGM (from bregma, AP 3.1 mm, ML 1.8 mm). Several injections of a mixture of *ssAAV-8/2-hSyn1-hM4D(Gi)_mCherry-WPRE-hGHp(A)* and *ssAAV-8/2-hSyn1-oChIEF_ChrimsonR_mCitrine-WPRE-SV40pA* (in a 1:4 viral particle ratio) were performed sequentially in different locations (AP 3.05 mm, ML 1.75 mm, DV 3.2 and 3.5 mm; second location AP 3.2 mm, ML 1.85 mm, DV 3.3 and 3.6 mm) so that a total of 4 injections of 0.5 μL were made to distribute the virus evenly through the desired location. Injections were made using a pressurized pico-spritzer and a pulled glass pipette. After viral injection, animals were kept for 4 weeks to ensure maximum virus expression through axon terminals.

For recording experiments, animals were anesthetized with 1.8 mg/kg urethane (i.p.), injected with 0.1 ml lidocaine (s.c.) in the incision points, and placed in a stereotaxic holder (Narishige, Japan). A trephine was drilled above the lateral amygdala (from bregma, AP 1.5 mm, ML 3.1 mm) and the 32 channel opto-electrode was lowered whilst stimulating until a maximum response was located (DV 4±0.2 mm). Body temperature was maintained constant at 37 °C through a feedback-regulated heating pad. Before recording, tissue was left for 10–20 mins for accommodation. Input-Output test curves were recorded for both intensity and pulse length 30 minutes before and 1 hour after drug injections using 1 ms-long pulses at 0.33 Hz (450 nm, 110 mA light intensity, aprox~14 mW, see Extended Fig. 8 Panel A). For electrical stimulation experiments (see Extended Fig. 8, Panels F-H), a stimulation bipolar twisted platinum-iridium microelectrode was placed in the internal capsule (AP −1.7 mm, ML 2.5 mm, DV 4.0 mm) to target the axons innervating the amygdala. Electrical pulses were given at an intensity that evoked 80% of the maximum response as biphasic 100 μs long pulses at 0.33 Hz.

Raw data was filtered (0.1 - 3000 Hz), amplified (100x), digitized and stored (10 kHz sampling rate) for offline analysis, with a tethered recording system (Multichannel Systems, Reutlingen, Germany). Analysis was performed using custom routines. After completion of the experiment, animals were sacrificed by decapitation and brains were extracted and kept in 4% paraformaldehyde overnight (RT). Brains were then sliced (50 μm) and inspected for viral infection location, extent, and electrode placement.

### Molecular modeling

The crystal structure of human muscarinic M4 receptor [Protein Data Bank (PDB) code: 5DSG] was used (after removing the bound ligand, tiotropium) for our modeling study. We made the Y113^3.33^G and A203^5.56^G in the M4 structure to create a DREADD (hM4Di). JHU37160 was prepared using LigPrep (Schrodinger LLC: New York, NY, 2017). The p*K*a calculations using Epik from Schrodinger suite (release 2017–4) and Chemicalize predicted that the piperazine nitrogen closer to the ethyl moiety has p*K*a values of 7.4–7.5 and we therefore protonated that nitrogen. Docking was performed using induced-fit docking protocol ^23^(Schrodinger LLC: New York, NY, 2017) with the OPLS3 force field. Four largest docking clusters with various orientations of the dibenzodiazepine moiety in the ligand binding site were identified using Clustering of Conformers in Maestro. The poses with the lowest docking scores from each of these four clusters were chosen for further MD simulations. Desmond MD systems (D. E. Shaw Research, New York, NY) with OPLS3 force field was used for the MD simulations. hM4Di was placed into explicit 1-palmitoyl-2-oleoyl-sn-glycero-3-phosphocholine lipid bilayer (POPC) using the orientation of the 5DSG structure from the Orientation of Proteins in Membranes database. Simple point charge (SPC) water model was used to solvate the system, charges were neutralized, and 0.15 M NaCl was added. The total system size was ~96000 atoms. The NPγT ensemble was used with constant temperature (310 K) maintained with Langevin dynamics, 1 atm constant pressure achieved with the hybrid Nose-Hoover Langevin piston method on an anisotropic flexible periodic cell, and a constant surface tension (x-y plane). The system was initially minimized and equilibrated with restraints on the ligand heavy atoms and protein backbone atoms, followed by production runs at 310 K with all atoms unrestrained. Two independent trajectories for each of the four JHU37160 poses were collected with an aggregated simulation length of 10.56 μs. Only one of the four poses both retained its ionic interaction with Asp112^3.32^ and showed convergence between the two trajectories, and thus was chosen for our further analysis shown in Fig. 3M.

### Statistics

Sample sizes were chosen based on our results from previous experiments. Depending on experiment, we used paired/two-sample t-tests or single factor and multifactor ANOVAs with *Dunnett’s or Tukey post-hoc tests*, taking repeated measures into account where appropriate. All statistical tests were evaluated at the *P*≤0.05 level and main figure statistics are shown in Table S1.

## Supporting information

## Acknowledgements

This work was supported by the NIDA (ZIA000069), NIMH (ZIAMH002793, ZIAMH002795), NIA, and NINDS Intramural Research Programs, the NIA (R21AG054802) to A.G.H, the NIBIB (P41EB024495) to M.G.P and by a European Research Council (ERC) 679714 STC grant to S.N. and Lundbeck Foundation fellowship R264-2017-3189 to A.M. We thank NIMH’s Molecular Imaging Branch (Chief, Robert Innis) for imaging the macaques with intracerebral injection of DREADDs. We thank Marisela Morales, Amy Newman, and Sergi Ferré for access to instrumentation, Hirsch Davis for access to pharmacological screening resources and Jian Jin for providing C13, C21, and C22. We thank Robert Dannals, Polina Sysa-Shah, James Engles, Nancy Ator, Taek-Soo Lee, Ben Tsui, and Peter Koncz for access to resources and technical support. We also thank Alex Cummins for assistance with macaque tissue preparation, Walter Lerchner and Violette der Minassian for lentivirus production, J. Megan Fredericks, Janita Turchi and Jalene Shim for compound formulation and administration, and Michael Frankland for assistance with blood sampling. M.M. is a cofounder and owns stock in Metis Laboratories.

## Author Contributions

All coauthors reviewed the manuscript and provided comments. J.B., M.A.E., F.H., J.L.G., M.S.S., A.M.A, S.L, M.B., C.R., M.F., S.S.S., S.T., S.S.Z., R.L.G., A.M., I.M.G.F., N.A., and J.Y.L. performed experiments, chemical synthesis and/or analyzed data. J.B., M.A.E., F.H., J.L.G., M.S.S., A.M.A, A.M., J.Y.L, V.W.P, R.B.I., R.M., P.M., L.S., D.R.S., S.V.M., S.N., A.G.H., B.J.R., and M.M. designed and/or supervised experiments and syntheses. M.G.P. and A.B. provided access to resources and support. J.B. and M.M. wrote the paper with input from all authors. M.M. conceived the study.

