## Supplementary material for "Chemogenetic ligands for translational neurotheranostics"

### Extended data.

Jordi Bonaventura[1†], Mark A. Eldridge[2†], Feng Hu[3†], Juan L. Gomez[1], Marta Sanchez-Soto[4], Ara M. Abramyan[5], Sherry Lam[1], Matthew Boehm[1], Christina Ruiz[6], Mitchell Farrell[6], Andrea Moreno[7], Islam Mustafa Galal Faress[7], Niels Andersen[7], John Y. Lin[8], Ruin Moaddel[9], Patrick Morris[10], Lei Shi[5], David R. Sibley[4], Stephen V. Mahler[6], Sadegh Nabavi[7], Martin G. Pomper[3], Antonello Bonci[11], Andrew G. Horti[3]*, Barry J. Richmond[2]*, Michael Michaelides[1, 12]*

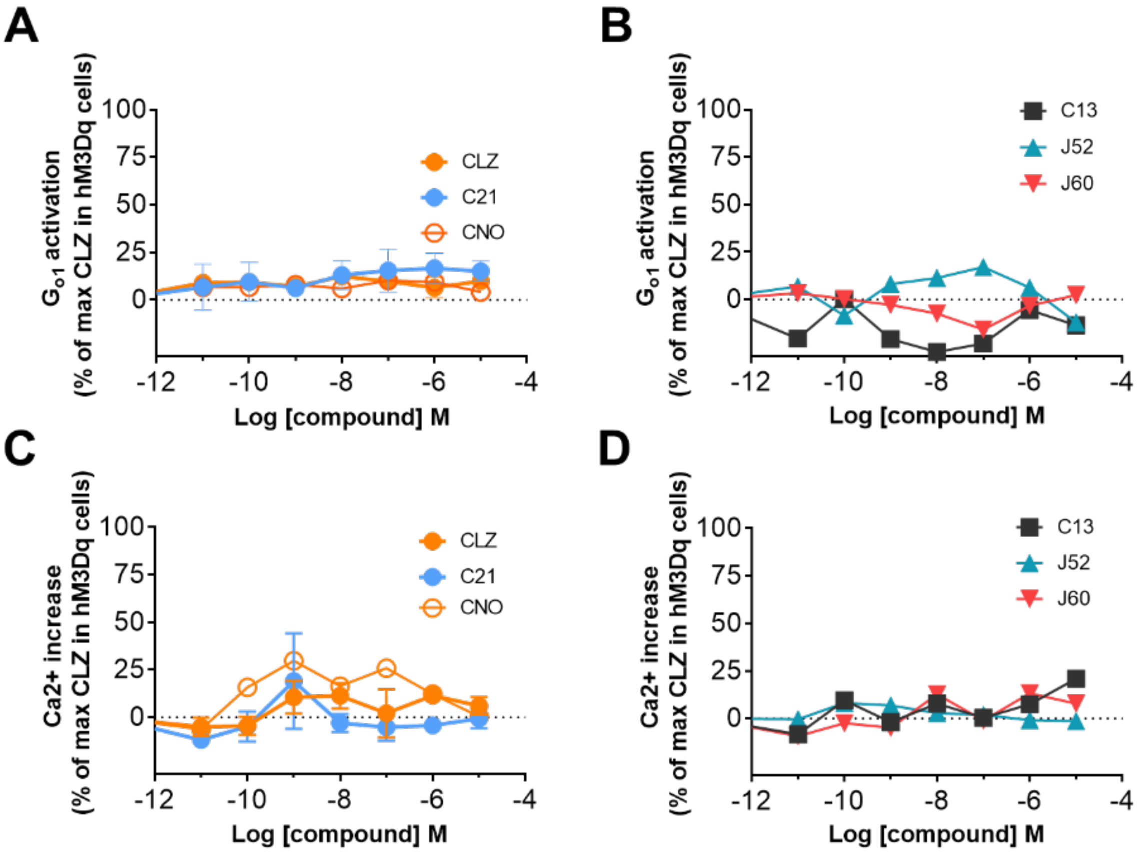


**Extended Figure 1. Negative controls for the cell-based functional assays. (A, B).** HEK-293 cells transfected with $G_o\alpha$-Rluc8, ß1 and γ2-mVenus but not DREADD did not show any functional response to Compound 21 (C21), clozapine (CLZ), CNO, Compound 13 (C13), JHU37152 (J52) or JHU37160 (J60). **(C, D).** HEK-293 cells transfected with GCaMP6f but not DREADD did not show any functional response to C21, CLZ, CNO, C13, J52 or J60. Data is shown as Mean ± SEM.

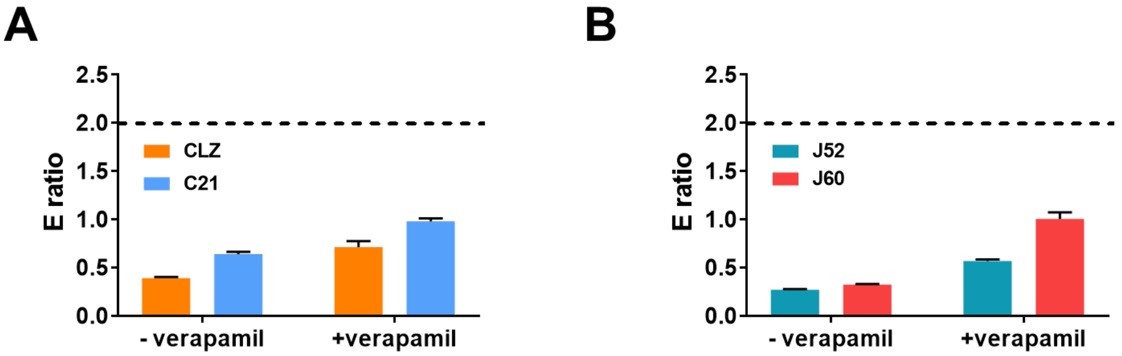


**Extended Figure 2. P-glycoprotein (P-gp) efflux assay. (A, B).** Efflux ratios for clozapine (CLZ), C21, JHU37152 (J52) and JHU37160 (J60) in the P-gp assay. If a compound has an efflux ratio greater than two in the absence of verapamil, and verapamil reduces the efflux ratio, it indicates that the compound is a substrate of P-gp. Hence, none of these compounds is a substrate for P-gp. Data is shown as Mean ± SEM.

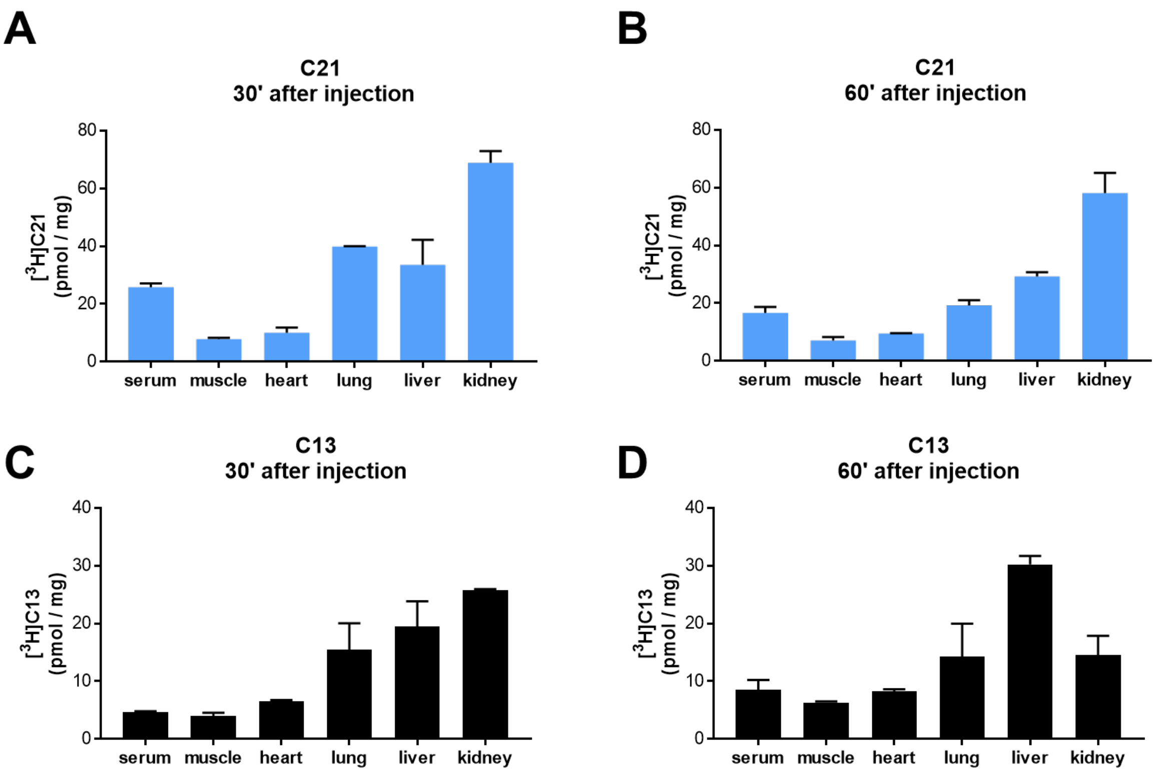


**Extended Figure 3. Biodistribution of C21 and C13. (A, B).** Concentration of [$^3$H]Compound 21 (C21) in major organs 30 or 60 min after an intraperitoneal injection of 2 μCi/g of C21. (**C, D).** Concentration of [$^3$H]Compound 13 (C13) in major organs 30 or 60 min after an intraperitoneal injection of 2 μCi/g of C21 or C13. Data is shown as Mean ± SEM.

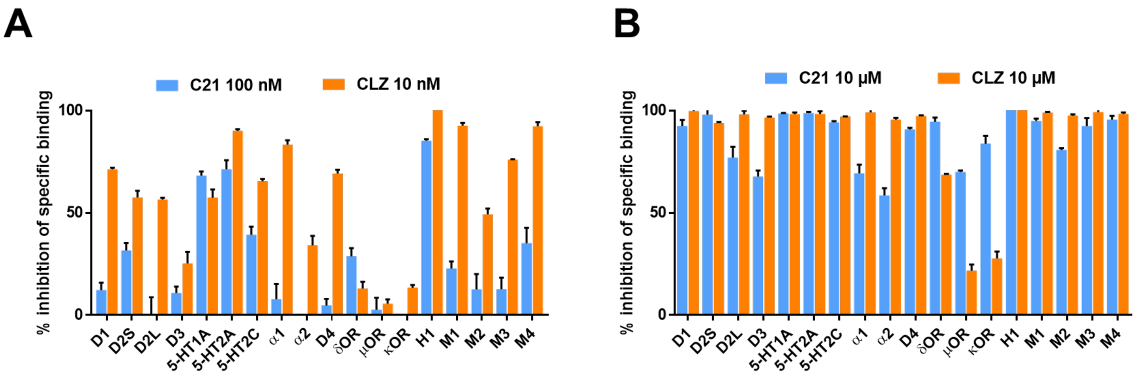


**C**

| | Ki values (nM) | | |
|---|---|---|---|
| | CLZ | J52 | J60 |
| 5-HT1A | 93.5 | ND | ND |
| 5-HT2A | 3.1 | 18.2 | 8.1 |
| 5-HT2C | 42.3 | ND | 31.9 |
| D1 | ND | ND | 23.5 |
| D2S | 7.9 | 7.8 | 4.9 |
| D4.4 | ND | ND | 14 |
| H1 | 1.1 | 4.6 | 1.4 |
| M1 | 8.3 | 6.2 | 3.1 |
| M2 | 85.1 | ND | 42.6 |
| M3 | 30.4 | 21.9 | 24.3 |
| M4 | 6.6 | 5.7 | 3.4 |

**Extended Figure 4. Endogenous binding screens.** (**A**) Inhibition of specific binding at endogenous potential targets by Compound 21 (C21) or clozapine (CLZ). CLZ has an affinity of ~2 nM whereas C21 has an affinity of ~200 nM and at equipotent concentrations they show similar affinity profiles for endogenous receptors. (**B**) At higher concentrations (10 µM) they show specific binding to multiple targets. Importantly, C21 binds to opioid receptors but clozapine does not. (**C**) Ki values for CLZ, JHU27152 (J52) or JHU37160 (J60) for receptors at which a 100 nM concentration produced at least 50% of inhibition of the selective ligand specific binding. The three compounds show similar binding profiles at dopamine and muscarinic receptors but not at serotonin receptors. Given that clozapine acts as an antagonist or weak partial agonist at these targets, and that cell-based functional assays in HEK-293 cells endogenously expressing those receptors did not show any response to J52 and J60, (Fig S1) it is likely that J52 and J60 will behave as antagonists for the endogenous targets whereas they are potent agonists for DREADDs. data is shown as Mean ± SEM.

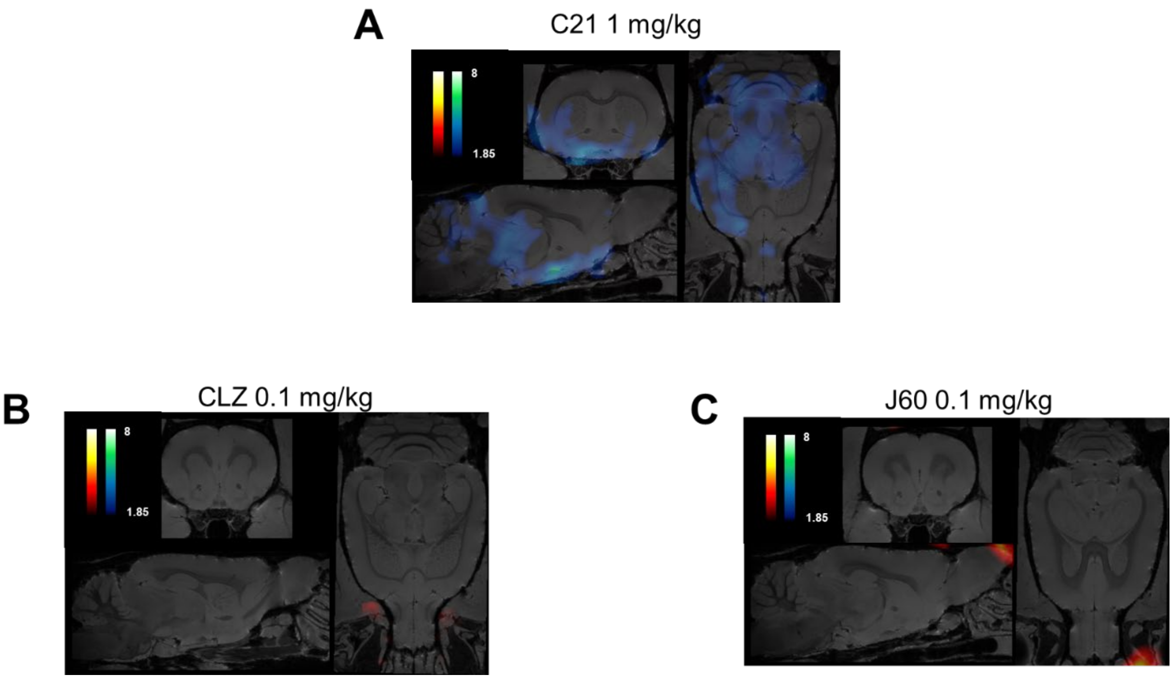


**Extended Figure 5. Patterns of metabolic activation in WT mice. (A)** Whole brain $^{18}$F-FDG metabolic mapping in WT mice shows decreased metabolic activity of multiple networks after a systemic administration of Compound 21 at 1 mg/kg. (**B, C**) Equipotent doses (0.1 mg/kg) of clozapine (CLZ) or JHU37160 (J60) did not show any significant changes in brain metabolic activity.

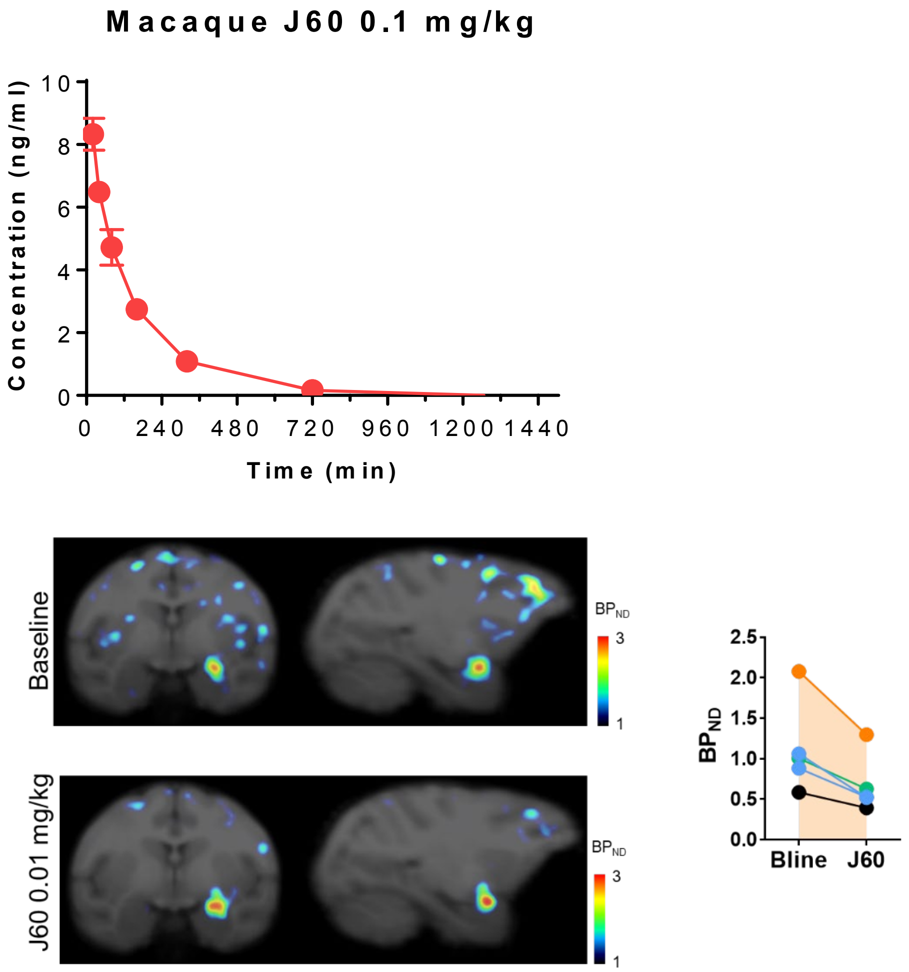


**Extended Figure 6. JHU37160 bioanalytics and target engagement in monkeys.** (**Top**) JHU37160 concentration in rhesus macaque serum after a 0.1 mg/kg systemic dose. (**Bottom**) J60 at 0.01 mg/kg systemic dose blocks [$^{11}$C]clozapine binding to hM4Di in monkeys. Data is shown as Mean ± SEM or individual values.

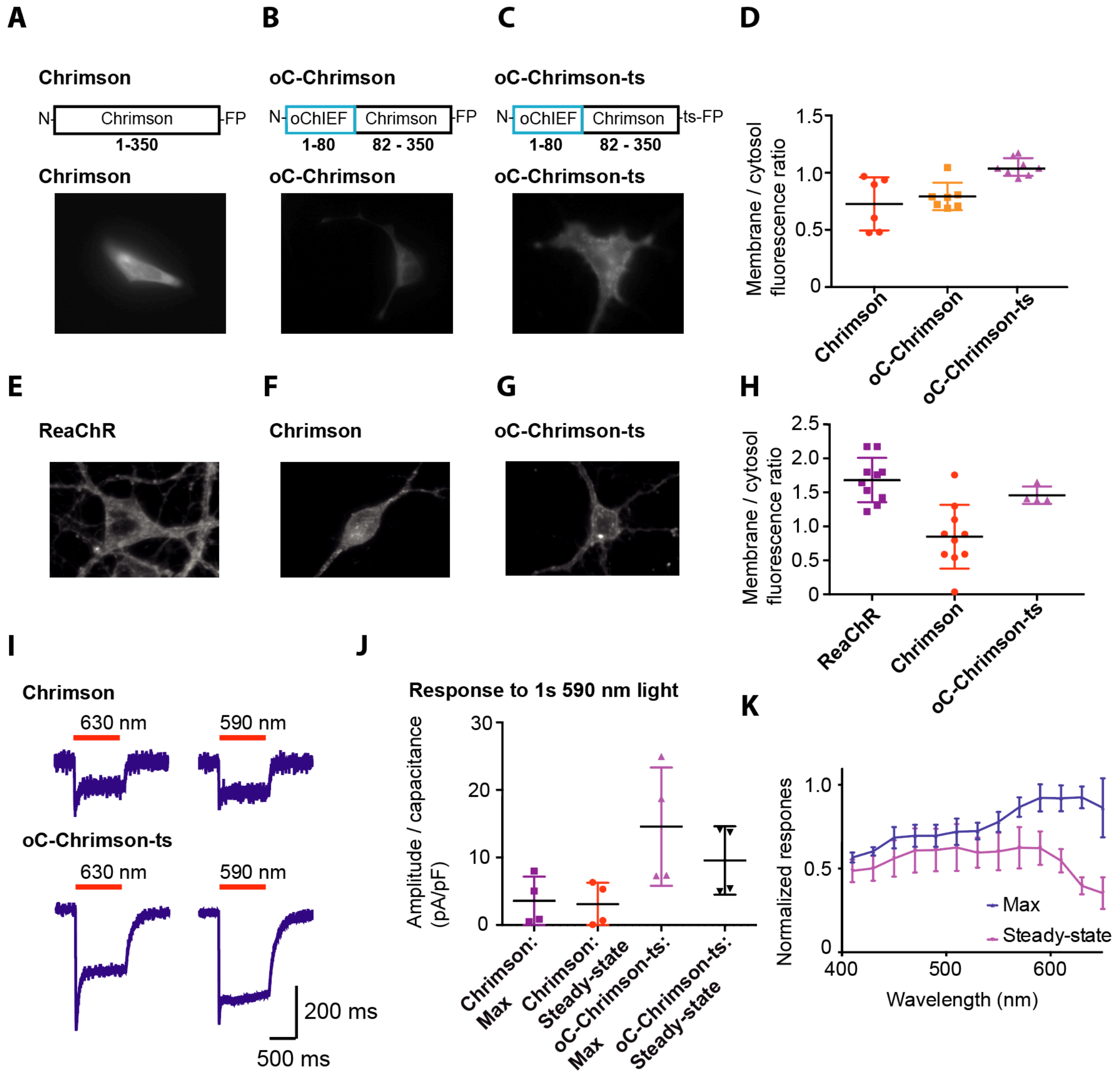


**Extended Figure 7. Chrimson variant generation and characterization.** (**A**) The expression of Chrimson-FP construct in HEK293 cells show mostly cytosolic expression pattern. (**B**) oC-Chrimson is a chimeric protein of N-terminus of oChIEF (residues 1-80) and Chrimson (residues 82-350) has similar cytosolic expression pattern to Chrimson in HEK293 cells. The N-terminus of oChIEF was used as it contained a predicted signalling peptide sequence for membrane exporting and the utilization of the same N-terminus sequence in ReaChR and oChIEF resulted in superior membrane trafficking. (**C**) oC-Chrimson-ts had additional trafficking signal (amino acid sequence, KSRITSEGEYIPLDQIDINV) placed between Chrimson and FP that resulted in membrane trafficking with minimal cytosolic aggregation in HEK 293 cells. (**D**) oC-Chrimson-ts has the best membrane to cytosolic fluorescence ratio compared to Chrimson and oC-Chrimson in HEK293 cells. (**E-G**) The expression of ReaChR, Chrimson and oC-Chrimson-ts in cultured cortical neurons. ReaChR and oC-Chrimson-ts show comparable membrane expression with Chrimson expression mostly within the cytosol. (**H**) The quantification of membrane to cytosol fluorescence ratio in cultured neurons. (**I**) Example traces of voltage-clamp recording of 1 s light evoked Chrimson and oC-Chrimson-ts photocurrent in HEK293 cells at indicated wavelengths. In most cells, the Chrimson response is below 100 pA whereas the oC-Chrimson-ts photocurrents are typically greater (>200pA) when cells of comparable fluorescence are tested. (**J**) Quantification of the maximum and steady-state (measured at 950 – 1000 ms) photocurrent of Chrimson and oC-Chrimson-ts to 1 s 590 nm light. (**K**) The normalized photocurrent response of oC-Chrimson-ts from 650 nm to 400 nm (n = 3). Similar spectral response for Chrimson cannot be measured accurately due to the small size of photocurrent in most cells recorded. All graphs are shown as mean ± SD.

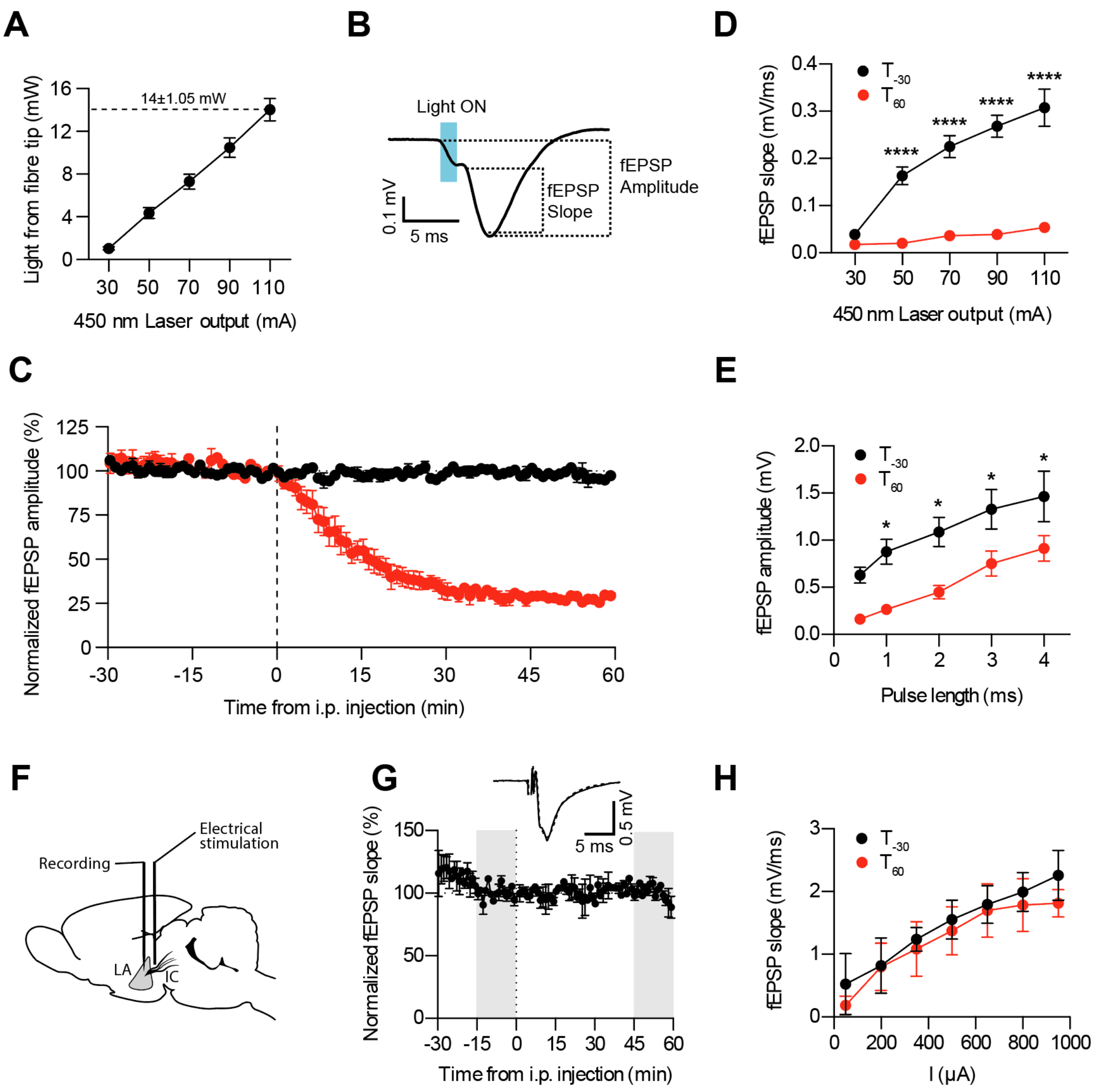


**Extended Figure 8.** ***In vivo*** **electrophysiological characterization of JHU37160. (A)** Corresponding measurements of light intensity stimulation and measured light output from the tip of the optic fibre (n=8), showing a consistent light delivery through experiments. (**B)** Field EPSP measurements of slope and amplitude on a representative waveform. (**C)** fEPSP amplitude measurements corresponding to the same experiments presented in Figure3O, P (n=4). (**D)** Input-Output test curves for light intensity 30 min before ($T_{-30}$) and 60 min after ($T_{60}$) JHU37160 injection (n=4). 2-way ANOVA effect of power F (4, 30) = 21.09 and time F (1, 30) = 210.0, post-hoc Sidak's multiple comparisons test shown as ****, p<0.0001 between T-30 and T60. (**E)** Input-Output test curves for pulse length 30 min before ($T_{-30}$) and 60 min after ($T_{60}$) JHU37160 injection (n=4). 2-way repeated measures ANOVA effect of pulse length F (4, 15) = 8.751 and time F (1, 15) = 47.06, post-hoc Sidak's multiple comparisons test shown as *, p<0.05 between T-30 and

T60. **(F)** Design of a control experiment with electrical stimulation of the Internal Capsule (IC) to LA pathway. (**G)** Time course of the effect of JHU37160 on non-hM4Di animals (n=2). Insert: representative waveforms from two selected time points (grey highlight in the graph) before and after infusion. (**H)** Input-Output test curves for stimulation intensity 30 min before ($T_{-30}$) and 60 min after ($T_{60}$) JHU37160 injection (n=2). 2-way repeated measures ANOVA effect of stimulation intensity $F (6, 7) = 3.101$ and time $F (1, 7) = 8.445$, post-hoc Sidak's multiple comparisons test not significant for all intensities between T-30 and T60. In all panels, data is shown as Mean ± SEM.

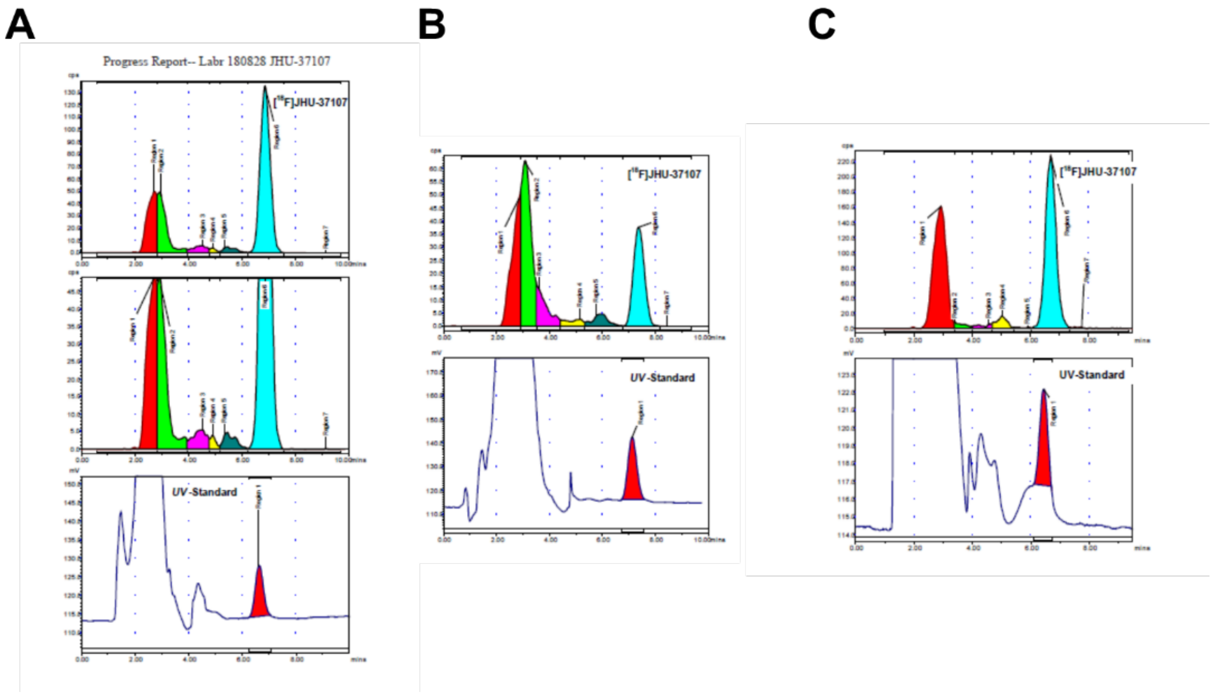


**Extended Figure 9. Radiometabolic analysis of [$^{18}$F]JHU37107. (A)** Radiochromatographic profile of monkey plasma, at 30 min after the IV administration of [$^{18}$F]JHU-37107 (J07). The parent radioactivity composition of plasma, region 6, was 56%. **(B)** Radiochromatographic profile of monkey plasma, at 180 min after the IV administration of J07. The parent radioactivity composition of plasma, region 6, was 24.2%. **(C)** Radiochromatographic profile of in vitro monkey whole blood (STD A), at 30 min of incubation with J07 at room temperature. The parent radioactivity composition of plasma, region 6, was 49.7%.
